# Division of Synthetic Cells Using a Genomically Encoded One-Protein Divisome

**DOI:** 10.64898/2026.07.21.739846

**Authors:** Charu Sharma, Nikki Nafar, Jacob Kerssemakers, José Ángel Fernández, Gijsje H. Koenderink, Cees Dekker

## Abstract

Synthetic biology is emerging as a powerful tool to understand life by building minimal synthetic cells whose functions are encoded within their genome. An essential function of cells is abscission, the final step of cell division that breaks the last connection between two emerging daughter cells. Abscission is challenging to reconstitute as it is energetically unfavorable to rearrange the cell membrane. Here, we demonstrate a minimal one-component abscission machinery for synthetic cells using the bacterial protein dynamin A (DynA), which is uniquely capable of driving scission inside membrane necks. Cell-free expression of this synthetic cell module is complicated by the large size of DynA (137 kDa) and the presence of a hydrophobic lipid-binding loop within its structure. We overcome these challenges by expressing DynA inside lipid vesicles with a high fraction of negatively charged lipids. We find that DynA enriches at the bridges of dumbbell-shaped vesicles, where it drives membrane scission and full division. The ability to monitor DynA dynamics through genomic encoding further provides hints about the mechanism of abscission including cooperative enrichment at the necks and constriction of the necks. Finally, we demonstrate that the smaller (71 kDa) dynamin D1 domain alone exerts the same function, which is beneficial for future integration with other synthetic cell modules since its much smaller gene size reduces the burden on the synthetic genome. This autonomously operating abscission machinery presents the first example of a self-dividing synthetic cell, marking an important step towards constructing a synthetic cell that is able to sustainably replicate.

## Introduction

Synthetic biology is emerging as a powerful tool to understand life by building synthetic cells from well-defined biomolecular components. A minimal synthetic cell is envisioned as a biomolecule-filled vesicle featuring essential cellular functions such as metabolism,^1,2^ growth,^3–5^ DNA replication,^5–7^ and division.^8–11^ A crucial requirement for a synthetic cell to be considered alive is that these functions must operate autonomously, where all the protein components driving the functions are encoded within a minimal synthetic cell genome. Various functions were so far been genomically encoded, including DNA replication,^5–7^ membrane growth,^3,5^ and membrane constriction from a spherical into a dumbbell shape,^12,13^, but not abscission. Abscission is the final step of division involving the cleavage of the intracellular neck that connects two newly formed daughter cells, separating them into two distinct individuals. It involves the local application of significant curvature stress to deform the cell membrane into a narrow neck that finally leads to scission. Abscission is very challenging to reconstitute, as it requires overcoming the membrane bending energy and line tension within the neck, to constrict it to a critical neck radius of <3 nm, beyond which the inner leaflets may fuse and form a stalk-like hemi-scission state that subsequently progresses towards full scission.^14,15^ As these topological transformations require the bilayer to deviate from its preferred planar configuration, abscission is energetically unfavorable. Cells have therefore evolved protein-based abscission machineries to manage this process.

In eukaryotes, abscission is performed by a complex ESCRT-III machinery,^16^ while archaea use a Cdv machinery.^17^ These multicomponent divisomes involve at least 3-4 proteins. The ESCRT-III abscission machinery has been reconstituted from purified components within giant unilamellar vesicles (GUVs).^18^ Notably, such multicomponent systems demand defined stoichiometries and precise spatiotemporal coordination to execute scission, making them highly challenging to adapt, for building a minimal autonomous abscission machinery for synthetic cells. This is where the Dynamin superfamily presents a compelling opportunity. Most members of the Dynamin family are one- or two- component systems, involved in membrane remodeling processes such as endocytosis and organelle division.^19^ Since these processes are topologically similar to abscission, Dynamins make highly attractive candidates for synthetic cell division. Unlike most Dynamin family members, which induce scission from outside the membrane, DynA from *Bacillus subtilis* (*B. Subtilis*) provides an interesting exception, as it operates in reverse topology, i.e., from within the cell. Strikingly, it is a one-component system in which the functionalities of its two subunits, D1 and D2, are combined within a single fusion protein. DynA is involved in membrane repair and defense and has also been suggested to play a role in *B. subtilis* cell-division. When DynA and the lipid-raft organizing protein floT were deleted from *B. subtilis*, the bacteria became highly elongated, filamented, and irregularly shaped, a classic hallmark of severely impaired cytokinesis.^20^ Moreover, DynA has been found to colocalize with bacterial ring-forming FtsZ, suggesting that DynA in bacteria plays a role in cell-division.^21^ All these factors make DynA a particularly suitable candidate for synthetic cell abscission.

In vitro, purified DynA was recently found to drive abscission of pre-deformed dumbbell-shaped giant vesicles.^10^ However, DynA has yet to be genomically encoded, to enable to exploit its full potential as an autonomously operating minimal abscission machinery for synthetic cells. Although DynA is a one-component division machinery, it is a relatively large protein (137 kDa), making it challenging to express using cell-free systems, which often have a limited capacity to synthesize full-length proteins larger than 100 kDa.^22^ Even if full-length synthesis is achieved, the hydrophobic membrane-binding domain can render the protein prone to aggregation. Moreover, the mechanism underlying DynA-induced abscission remains poorly understood, which further impedes its application as a cell-division machinery. Genomic encoding of DynA will not only enable its autonomous operation but may also provide temporal resolution of the abscission process, from DynA expression and localization at membrane necks to the abscission event, something that cannot be achieved by reconstitution of the purified protein.

Here, we show that a one-component DynA machinery can be autonomously produced from DNA by cell-free expression within dumbbell-shaped GUVs, where it localizes to the necks and drives autonomous scission (Figure 1a). We genetically fused DynA to a fluorescent protein to track the live expression and enrichment of DynA via fluorescence microscopy. We found that cell-free expressed DynA was functionally active only when expressed in giant vesicles prepared with high percentages of negatively charged lipids in their membranes. Time-dependent imaging tracking of the initial mechanistic steps towards abscission showed cooperative enrichment at the necks and constriction of the necks. We show that, right after the DynA induced division, the newly generated daughter cells are held together by weak van der Waals forces and can be physically separated from each by applying a weak (∼1.4 nN) force. Finally, we find that D1, a truncated part of DynA, is similarly effective autonomous abscission machinery for synthetic cells, which presents significant advantages as its much smaller size (71.2 kDa) facilitates faster, more-complete, and higher-yield expression in synthetic cells. Overall, the findings present the first example of a self-dividing synthetic cell driven by a one-protein divisome.

**Figure 1.**
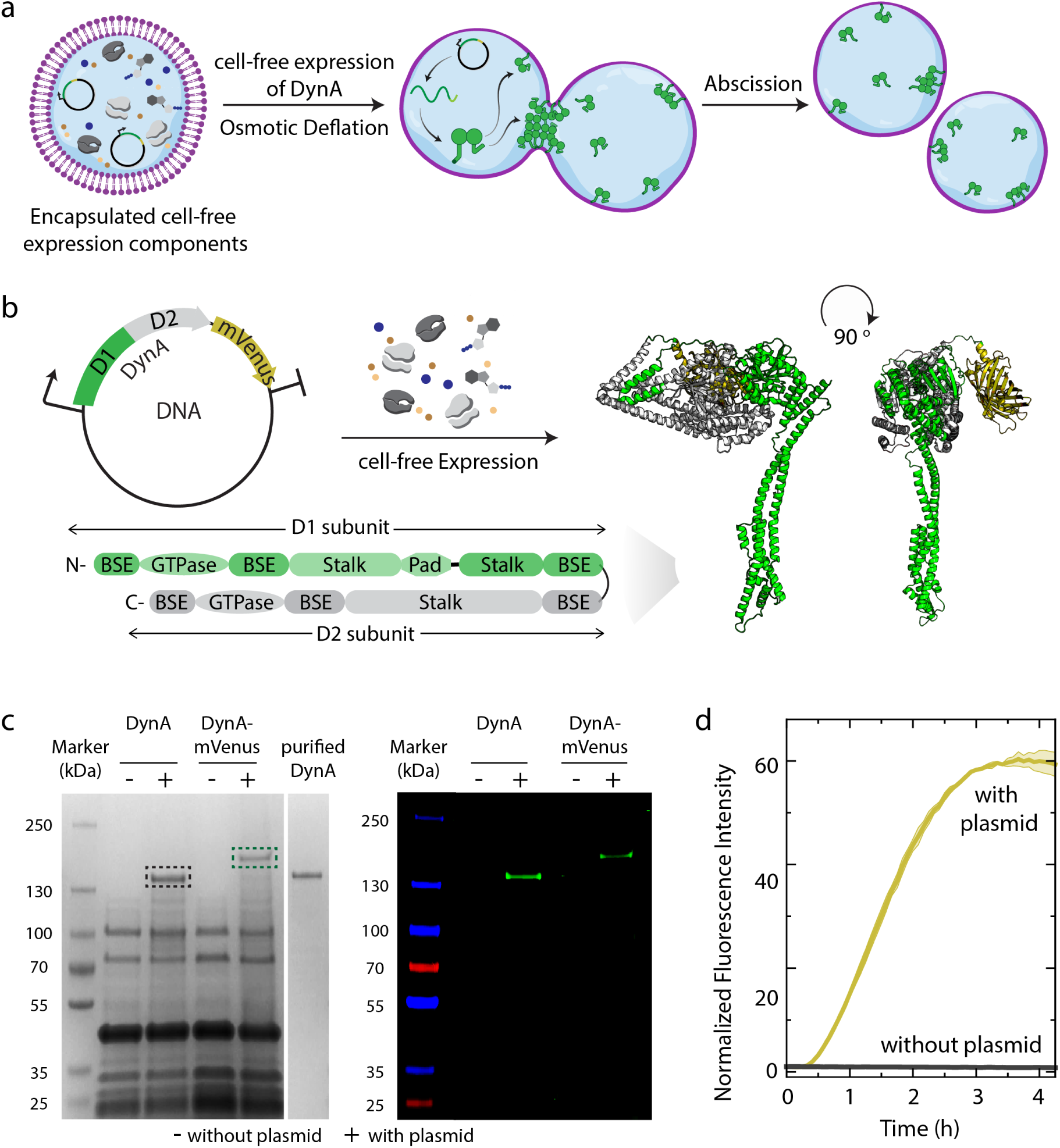
Genomically encoding one-component DynA machinery for bottom-up synthetic cell division. **a)** Schematic depicting the reconstitution of a minimal division system for synthetic cells. Cell-free expression components along with a DNA template encoding DynA are encapsulated inside pre-deformed lipid vesicles. In vitro expressed DynA localizes at the neck of the dumbbell where it divides the dumbbell into two separate vesicles by membrane abscission. **b)** Schematic illustration of in vitro expression of DynA-mVenus using PURE cell-free expression system. DynA is fused with mVenus at the C-terminal causing no structural differences as confirmed with AlphaFold3 predictions. Bottom part of figure represents the domain architecture of DynA predicted via structural modeling where BSE and Pad stand for bundle signaling element and paddle like domain, respectively. **c)** Translated products were analyzed by SDS-PAGE. Left: post-stained with Coomassie stain. Right: In vitro expression was carried out in the presence of tRNA_Lys_ preloaded with a BODIPY-conjugated lysine (GreenLys) and the translated products were analyzed by fluorescence imaging of SDS-PAGE gel. 10 ng/uL of DynA or DynA-mVenus template was used. **d)** Time-resolved normalized fluorescence of DynA-mVenus during cell-free expression. The fluorescence intensity at time, t is normalized with respect to baseline intensity at t =0. The results represent an average contribution from two independent measurements; the shaded region depicts the standard deviation (SD).

## Results

### Full length Dynamin A can be expressed using a cell-free system

Encapsulation of a genome and its expression machinery encoding desired cellular functions within a compartment such as a liposome is a compelling strategy for recapitulating life-like functions in synthetic cells. A cell-free expression system comprising transcription and translation components acts as a simplified cytoplasm to produce proteins from DNA templates in vitro.^23^ However, synthesizing larger proteins (>100 kDa) using cell-free expression systems often suffers from low ribosome recycling efficiency and premature translation termination, resulting in low protein yields and the accumulation of truncated polypeptides.^22^

DynA in *B. subtilis* is thought to have arisen through the duplication and subsequent fusion of an ancestral dynamin-like gene, resulting in a single, but large (137 kDa) protein containing two connected dynamin-like domains, D1 and D2. The D1 subunit of DynA is crucial for membrane tethering and fusion, while the D2 subunit facilitates membrane fusion via unknown mechanism.^24^ Here, we first checked whether full-length DynA can be successfully synthesized using a cell-free expression system. We used PUREfrex2.0, a reconstituted *E. coli* cell-free expression platform chosen for its low protease/nuclease activity, high controllability, and ability to produce functional membrane-binding proteins.^25^ Aware of the anticipated challenges associated with the cell-free expression of a large protein such as DynA, we first performed several preliminary checks on the DNA template. Specifically, we assessed codon usage and other sequence features to minimize the likelihood of premature truncation arising from template-related issues and to ensure that the DynA-encoding DNA template was optimized for cell-free expression. Specifically, we confirmed a low CG content (39%) in the first 6 codons of the open reading frame, the absence of significant secondary mRNA structure around the start codon, and the absence of any proline repeats in the entire open reading frame.^23^ To build a DNA template, the sequence encoding wild-type DynA was fused at its C-terminus via a flexible linker to a fluorescent protein (mVenus) for visualization. We chose mVenus, due to its higher extinction coefficient, improved quantum yield, and monomeric behavior compared to GFP previously used to tag DynA in cells (Figure 1b).^26,27^ AlphaFold3 did not indicate any structural differences by the addition of mVenus to DynA (Figure S1).

Succesful expression of both DynA and DynA-mVenus in PURE was established, as confirmed by SDS-PAGE, with Coomassie staining revealing bands at the expected molecular weights (137 kDa and 164 kDa, respectively), alongside background PURE proteins (Figure 1c, left panel). Quantification indicated the production of ∼1.1 µM DynA using a plasmid concentration of 10 ng/uL, well exceeding the levels (250 nM) required for abscission (Figure S2).^10^ To test whether any truncated products were present, expression was carried out in the presence of fluorescently labeled lysine residues, which co-translationally incorporate within the expressing proteins rendering them fluorescent for visualization by fluorescent gel imaging (Figure 1c, right panel). Only bands corresponding to full-length DynA and DynA-mVenus were observed, without any truncated polypeptides. Finally, correct folding of DynA-mVenus was validated through its intrinsic fluorescence, which depends on chromophore maturation:^28^ Time-resolved measurements showed a 60-fold fluorescence increase compared to baseline during expression, reaching steady-state within 3.3 hours (Figure 1d, Figure S3).

### Negatively charged DOPG lipids promote expression of functionally active Dynamin A

After establishing successful cell-free expression and proper folding of full-length DynA-mVenus in bulk, we next assessed whether DynA-mVenus is expressed and functionally active in confinement within GUVs as a cell-sized membrane system. GUVs were prepared from a lipid mixture of DOPC, DOPE-PEG2000, DOPG, and Cy5-DOPE at a molar ratio of 89.75:2:8:0.25. DOPC was selected as the main component due to its low phase transition temperature (fluid at room temperature) and zwitterionic nature,^29^ which support thermodynamically stable GUVs in high yields. The membranes were doped with 8 mol% negatively charged DOPG to facilitate DynA binding.^10,26^ Additionally, 2 mol% DOPE-PEG2000 was included as its flexible PEG chain enhances the vesicle yield and reduces non-specific protein adsorption at membranes, thereby limiting vesicle aggregation. We used the Inverse-Emulsion method to produce GUVs because of its high encapsulation efficiency across a wide range of macromolecules.^30–32^ First an Inner Aqueous Solution (IAS) containing PURE components (buffer, enzymes, ribosomes), 20 ng/µL of DynA-mVenus template, and 200 mM sucrose was emulsified in a lipid-in-oil phase to form water-in-oil droplets. This emulsion was then layered onto an outer aqueous solution (OAS) containing Home-made PURE Solution I and 200 mM glucose. Upon centrifugation, droplets traversed the oil-water interface to form GUVs encapsulating the cell-free expression system, which were subsequently incubated at 37 °C. Sucrose and glucose are widely used in GUV preparation to control osmotic conditions and are compatible with PURE encapsulation.^31,33^ By using an asymmetric distribution of sucrose and glucose, we were able to generate dumbbell-shaped GUVs making use of the positive spontaneous curvature at the membrane (∼1 µm^-1^) generated by the sugar asymmetry^34^. Under near isotonic conditions (osmolality ratio between OAS and IAS is 0.9), we obtained 41% dumbbell-shaped GUVs, which act as an excellent model system for our in vitro cell-division studies (Figure S4).

DynA-mVenus was successfully expressed within GUVs, as confirmed by an increased fluorescence over 5 hours of incubation. However, the protein predominantly formed aggregates that increased in size over time (Figure 2a, left panel; Figure S5). Protein aggregation in cell-free systems can arise from multiple factors, including suboptimal environmental conditions (temperature, crowding, high protein concentration), presence of truncated polypeptides, absence of cellular folding machinery, and intrinsic protein properties such as exposed hydrophobic patches.^35^ These effects may be pronounced for DynA-mVenus given its large size (163 kDa) and its membrane-binding region.

**Figure 2.**
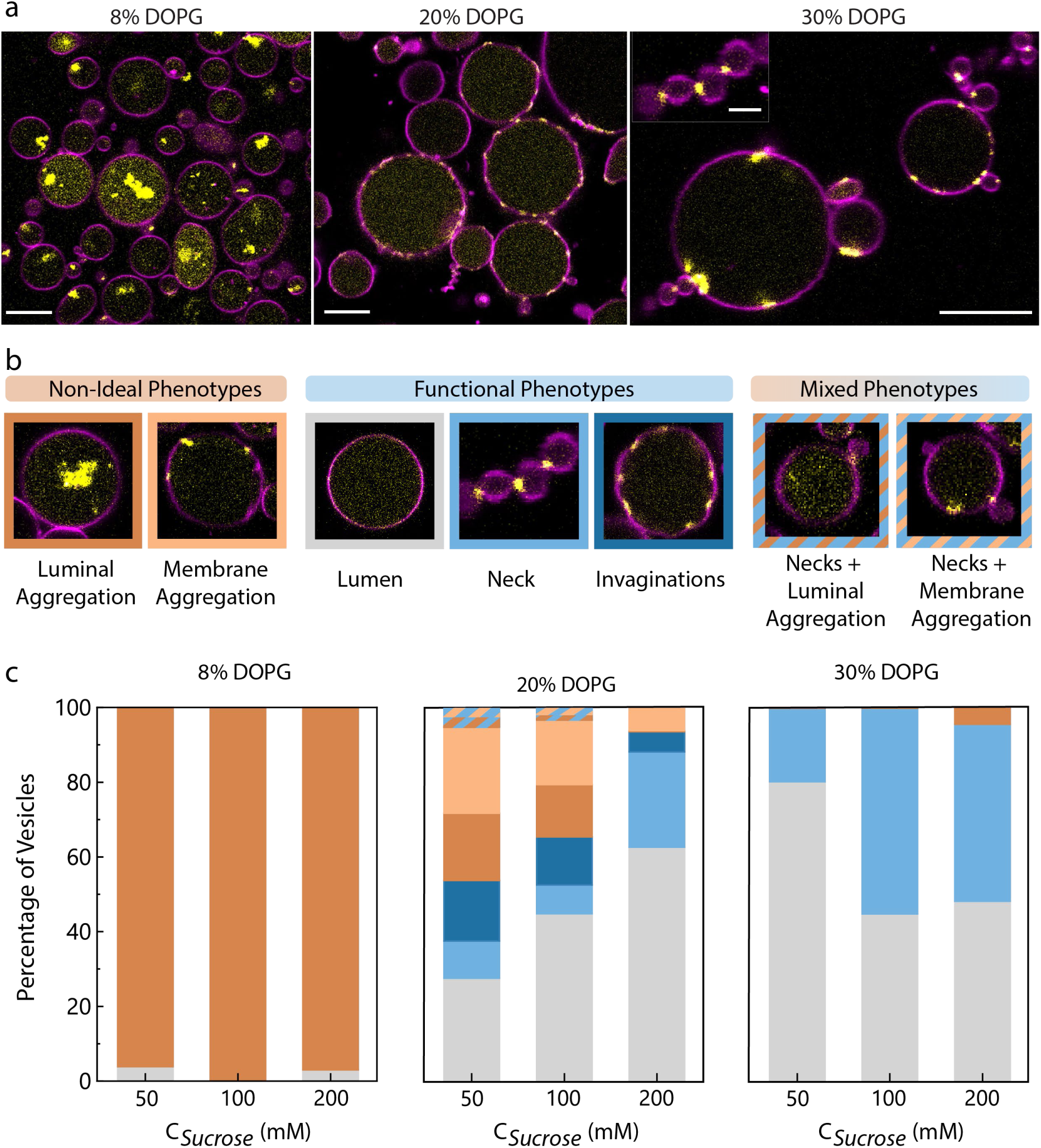
Membrane and lumen composition jointly affect in vitro expression of functionally active DynA. **a)** Representative images of in vitro expressed DynA-mVenus encapsulated within GUVs prepared with increasing DOPG concentrations (8%, 20%, and 30%) using Sucrose as a crowder (200 mM for 8% DOPG and 100 mM for 20% and 30% DOPG) and 20 ng/uL of DynA-mVenus template incubated at 37 °C. Scale bar: 10 µm, inset: 2 µm. **b)** Phenotypic classification of the vesicles based on state (aggregated or functional) and DynA-mVenus localization (luminal, curved sites or necks) into three categories: (i) Non-ideal phenotypes, consisting of aggregated DynA-mVenus localized either within the lumen or at the membrane, (ii) Functional phenotypes, with DynA-mVenus either remaining dissolved within the lumen or enriching at the necks or invaginated sites, (iii) Mixed phenotypes where DynA-mVenus is enriched at the necks in addition to showing luminal or membrane aggregation. **c)** Percentage distribution of vesicles representing 7 phenotypes of in vitro expressed DynA-mVenus within vesicles prepared with 8%, 20%, and 30% DOPG concentrations, where each DOPG condition is screened across three sucrose concentrations (50 mM, 100 mM, and 200 mM). 550-800 vesicles were phenotyped per condition. Color coding corresponds to the outlines in panel b. All images were taken after 4.5 hours of expression using 20 ng/µL DNA.

We first aimed to suppress aggregation by optimizing solution conditions, beginning with macromolecular crowders and folding chaperones. Given that protein folding normally occurs in a crowded intracellular environment, macromolecular crowders are often used in vitro to mimic such conditions. While crowders can enhance expression efficiency by increasing effective concentrations of transcription–translation components,^36^ excessive crowding can also promote aggregation.^37^ Sucrose can induce crowding-like effects via excluded-volume interactions.^38,39^ Therefore, we expressed DynA-mVenus at different sucrose concentrations, 50 mM, 100 mM, and 200 mM, and observed an 80% reduction in aggregation when sucrose was lowered from 200 mM to 100 mM (Figure S6). Reducing sucrose further to 50 mM provided only an additional 10% reduction. Therefore, all subsequent optimizations were performed with 100 mM sucrose. As the PURE system lacks chaperones for protein folding, we next introduced two well-characterized folding systems: DnaKJE and GroE, both known to enhance solubility of aggregation-prone proteins.^40,41^ DnaKJE acts co-translationally at the ribosome, whereas GroE encapsulates partially folded substrates within its cavity. DnaKJE and GroE reduced DynA-mVenus aggregation by 18% and 38%, respectively, but neither prevented full aggregation (Figure S7). Lowering the plasmid concentration from 20 ng/µL to 4 ng/µL further reduced aggregation by 24%, while decreasing the reaction temperature to 30 °C only limited the formation of larger aggregates (Figure S8). None of these solution-based conditions fully eliminated aggregation, so we turned our attention to the GUV lipids.

Cell-free expression of transmembrane proteins, which are often highly aggregation-prone, is routinely performed in the presence of DOPC liposomes, with optimization achieved by tuning lipid chain length or saturation, or by adding amphiphiles to promote membrane insertion and folding.^42^ This strategy is however uncommon for membrane peripheral proteins like DynA, which exist in both soluble and membrane-bound form and only transiently bind to the membrane surface. Accordingly, the role of membranes on cell-free expression of membrane-peripheral proteins has not previously been explored, largely because peripheral proteins do not insert into bilayers but instead interact via small hydrophobic motifs stabilized by electrostatic contacts through e.g. lysine or arginine residues in lipid-binding loops.^43^ Interestingly, a membrane-binding region of five amino acids was recently identified in the DynA D1 subunit, containing two lysines and three phenylalanines.^44^ Given this mixed electrostatic-hydrophobic interaction mode, we reasoned that the lipid composition of the membrane environment of DynA requires optimization. Because the lipid-binding loop of DynA is positively charged owing to the lysine residues, we increased the fraction of negatively charged lipids in the membrane. Given the ∼30% PG content of *B. subtilis* membranes,^24,45^ we first explored the binding of purified DynA to GUVs with increasingly negatively charged bilayers. Because higher DOPG fractions can reduce GUV yield during inverse emulsion preparation by impairing monolayer assembly and generating leaky, mechanically unstable vesicles,^46^ we restricted our analysis to membranes containing 8-30% DOPG, which spans the physiological PG content of *B. subtilis* membranes. Under these conditions, purified DynA exhibited preferential binding to membranes containing 20% and 30% DOPG, whereas it remained localized within the lumen of membranes containing 8% DOPG (Figure S9).

To test whether negative membrane charge plays a role in the correct folding of DynA-mVenus, we thus prepared GUVs in which the DOPG content was increased from 8% (used previously) to 20% and 30% (Figure 2a, middle and right panel). Strikingly, aggregation of the cell-free produced protein was markedly reduced at 20% DOPG and completely eliminated at 30% DOPG, directly supporting a role of the membrane charge in preventing DynA-mVenus aggregation. To quantitatively analyze the effect of DOPG concentration, we classified the spectrum of phenotypes observed (Figure 2b) into three categories: (i) non-ideal phenotypes, characterized by aggregated DynA-mVenus either in the lumen or at the membrane; (ii) fully functional phenotypes, in which aggregation was absent and DynA-mVenus remained soluble in the lumen or enriched at necks and invaginated regions, as seen previously for BAR proteins^47^ and curved DNA origami^48^; and (iii) mixed phenotypes, with characteristics of both category i and ii. We observed that protein aggregation was reduced by ∼66% in the 20% DOPG condition and prevented completely for 30% DOPG for expressions carried out in the presence of 100 mM sucrose (Figure 2c). Membrane neck localization was observed in 8% of 20% DOPG GUVs (Figure 2c, middle panel) and 55% of 30% DOPG GUVs (Figure 2c, right panel).

Overall, membrane composition thus emerged as a more effective parameter to express functional DynA-mVenus than other physical factors, such as temperature, plasmid concentration, or the addition of chaperones. To preserve high expression rates^49^ and maintain compatibility with other synthetic cell modules, the incubation temperature was kept at 37°C. Modulation of sucrose concentration further exhibited a DOPG-dependent effect, suggesting that the impact of sugar on protein aggregation is highly dependent on membrane lipid composition (Figure 2c). Notably, GUVs prepared with 30% DOPG and 100 mM sucrose yielded up to 55% of dumbbells with DynA-mVenus at necks, making it the most favorable condition for programming synthetic cell abscission.

### In vitro expressed Dynamin A senses membrane curvature

Performing DynA-mVenus expression in the optimized conditions in GUVs, we collected confocal z-stacks of GUVs over time during expression, enabling direct observation of DynA-membrane interactions. We identified three distinct behaviors of DynA-mVenus during its cell-free expression: In the first, and most frequent case, 55% of the GUVs showed phenotype of cell-free expressed DynA-mVenus formed clusters which localize at the highly curved necks of the pre-deformed GUVs (Figure 2c, Figure 3a). The extent of DynA-mVenus accumulation at the dumbbell necks was quantified using the enrichment ratio (E_R_) quantifying the ratio between the intensities at the neck and in the lumen, yielding an average E_R_ of 18.8 (Figure S10). We used the E_R_ as a quantitative criterion to distinguish GUVs exhibiting clear DynA clustering at the neck from those showing no detectable neck enrichment. These E_R_ values are comparable to those obtained with purified DynA (E_R_ = 56; Figure S11), where DynA was present at a concentration of approximately 18 µM within the clusters, corresponding to an estimated ∼1,300 DynA molecules per cluster. Furthermore, the enrichment ratio increased progressively as cell-free expression proceeded, indicating a time-dependent accumulation of DynA-mVenus at sites of high membrane curvature (Figure 3b).

**Figure 3.**
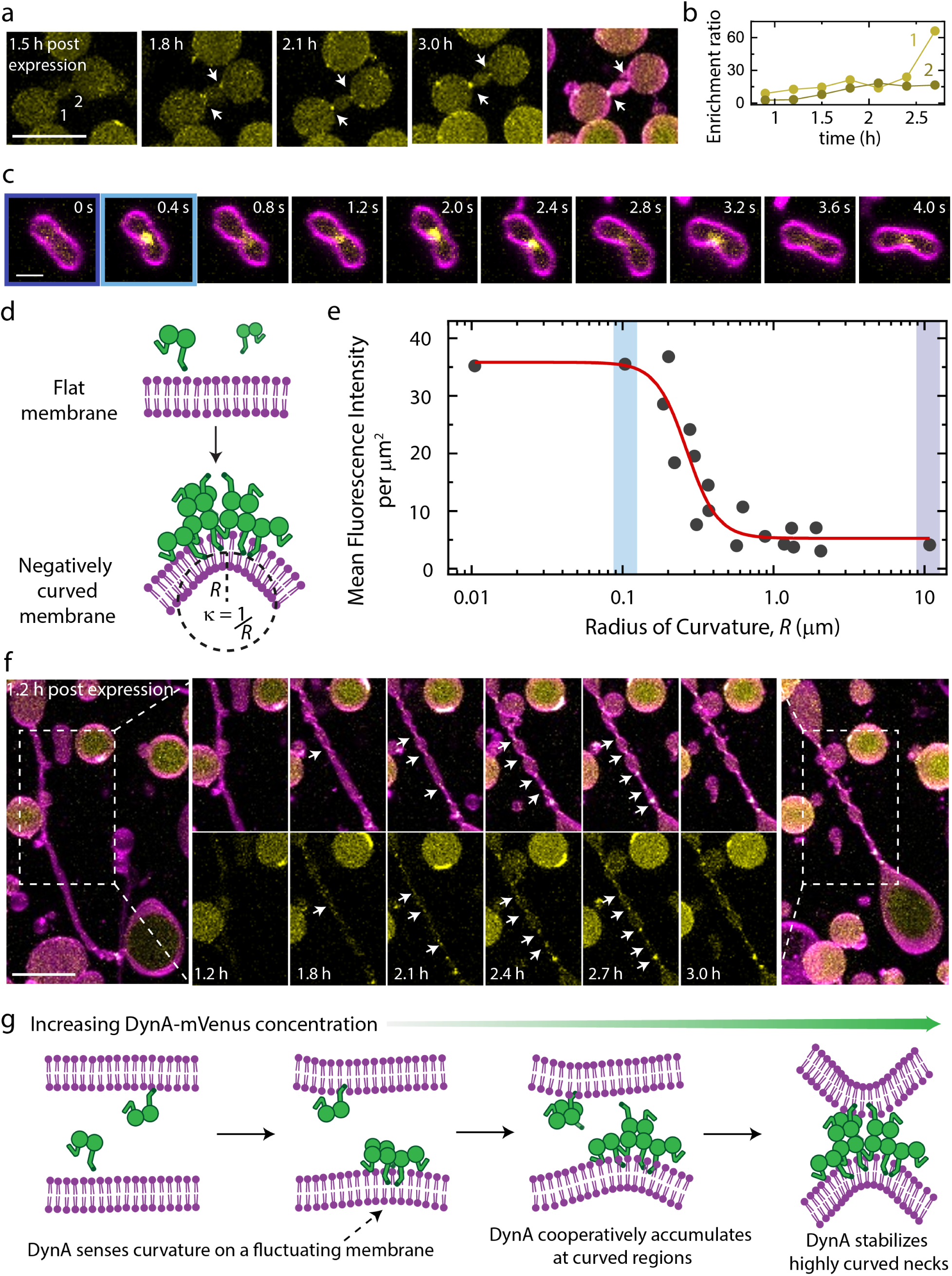
DynA cooperatively enriches at negatively curved sites and induces local spontaneous curvature. **a)** Time-resolved 3D projections of in vitro DynA–mVenus expression (20 ng/µL DNA, 100 mM sucrose, 30% DOPG) show preferential enrichment at the necks (indicated by arrows) of dumbbell-shaped vesicles. Scale bar: 20 µm**. b)** Enrichment ratio of DynA-mVenus at the necks of a dumbbell over 3 h of cell-free expression. **c)** Time series of a floppy GUV shows that in vitro expressed DynA-mVenus switches mainly between two states, either remaining homogeneously distributed within the lumen or forming clusters with varying fluorescence intensity that enrich at transiently generated negatively curved membrane sites. Scale bar: 2 µm. **d**) Schematic illustration of DynA-mVenus forming clusters exclusively at negatively curved sites, where the point curvature, κ, is determined by fitting a circle to the tip of the membrane deformation. The radius of this fitted circle, R, defines the radius of curvature and is the inverse of the point curvature. **e)** The radius of curvature (R) obtained from the time series in (c) is plotted against the mean fluorescence intensity of DynA–mVenus measured within a circular area of 1 µm² directly above the curvature measurement site. Fitting with a sigmoidal Hill equation yields a Hill coefficient of 4.6 (>1), indicating positive cooperativity. This suggests that DynA–mVenus remains largely unbound until R drops below ∼0.3 μm, whereas clustering always occurs and becomes insensitive for curvatures <0.1 μm. **f)** 3D projections of a pre-deformed vesicle show a wide cylindrical tube connecting two lobes transforming into a chain of pearls with multiple lobes, where each emerging neck becomes enriched with DynA-mVenus as expression progresses over 3 hours. The longitudinal contraction of the tube from 124 µm to 36 µm coincides with the time-dependent appearance of DynA-mVenus-localized necks, indicating that DynA stabilizes the constricted membrane topology and promotes further neck constriction. Scale bar: 20 µm. **g)** Proposed model for the constriction of highly curved necks by DynA, based on the experimental results in (a–f). At low DynA concentrations, the membrane tube remains in its original cylindrical shape. As DynA accumulates during cell-free expression, it senses transient negatively curved deformations arising from thermal fluctuations, where binding lowers the local membrane bending energy. With increasing DynA concentration, the protein cooperatively accumulates at these transiently curved sites. This establishes a positive feedback mechanism in which highly curved regions recruit more DynA, while the increasing local DynA concentration further reduces the energetic cost of neck formation, thereby stabilizing the constricted membrane topology and promoting further neck constriction.

In the second, less frequent, case, we observed GUVs with DynA-mVenus oscillating between two distinct states (Figure 3c), where the protein either spread homogeneously throughout the vesicle lumen, or it transiently formed clusters at curved sites (Figure 3c, d). To quantify this behavior, we correlated DynA-mVenus fluorescence with the local radius of curvature at the saddle-shaped neck (Figure 3e). This analysis revealed a clear transition between a homogeneous and clustered state of DynA-mVenus, with cluster formation sharply increasing once the radius fell below ∼0.3 μm, whereas clustering always occurred for curvatures <0.1 μm. A sigmoidal Hill equation to the data yielded a Hill coefficient^50^ of 4.6, indicating positive cooperativity. Notably, cluster formation was reversible, as clusters dispersed back into the diffuse state under relaxed GUV morphologies, highlighting a dynamic and curvature-dependent assembly and disassembly process.

Finally, in the third case, we observed GUVs connected by a wide cylindrical tube exhibiting membrane fluctuations (Figure S12a), which progressively deformed into a chain of pearls during expression (Figure 3f). Over ∼3 hours, the tube shown in Figure 3e underwent longitudinal contraction (from 124 µm to 36 µm; more examples in Figure S12) while generating multiple necks that were each enriched in DynA-mVenus. This pearling transition observed during progressive DynA expression can be explained in terms of the Helfrich model.^51,52^ At early stages, when the protein concentration was low, the membrane tube remained in its original cylindrical shape (Figure 3g). As DynA accumulated over time, it preferentially partitioned to regions of incipient neck curvature. The time-dependent emergence of multiple DynA-decorated necks, together with the cooperative binding of DynA to negatively curved membranes established in the second case (Figure 3e), suggests a positive feedback mechanism. Specifically, regions of high curvature recruit more DynA, while the increasing local concentration of DynA further reduces the energetic cost of neck formation, thereby stabilizing the constricted membrane topology and promoting further neck constriction.

### Dynamin A induces full scission into daughter cells that separate under a weak external force

Next, we asked whether neck-localized DynA clusters can drive complete scission of the neck to fully divide the vesicles. To test this, we probed the membrane connectivity between lobes using fluorescence recovery after photobleaching (FRAP) experiments.^10^ Here, fluorescently labelled lipids in one of the lobes of the dumbbell vesicle were photobleached, followed by measuring the fluorescence recovery of this lobe over time as lipids diffused across the neck region connecting the adjacent lobe. We quantified the degree of fluorescence recovery by the normalized intensity (N_i_), which measures the ratio between the final fluorescence intensity of the bleached lobe after recovery and that of a neighboring control lobe that was not photobleached. A final value of N_i_ = 1 indicated full recovery by free diffusion of lipids through the neck, hence signifying that no scission had occurred. In contrast, N_i_ = 0 corresponded to no recovery, indicating full scission with a loss of connectivity between the dumbbell lobes. An intermediate value of N_i_ reflected a partial recovery, consistent with hemi-scission where lipid diffusion restricted to the outer leaflet of the membrane.^10^ Consistent with previous experiments with purified DynA,^10^ we observed all three outcomes of FRAP experiments on chains of vesicles encapsulating cell-free expressed DynA-mVenus (Figure 4a). Pooling all FRAP data, we found that cell-free expressed DynA-mVenus induced membrane remodeling events in 42% of the GUVs, driving 27% of analyzed vesicles to full scission and 15% to hemi-scission (Figure 4b). These results establish DynA as a single-component synthetic cell division machinery that can be genomically encoded and functionally expressed under cell-free conditions.

**Figure 4.**
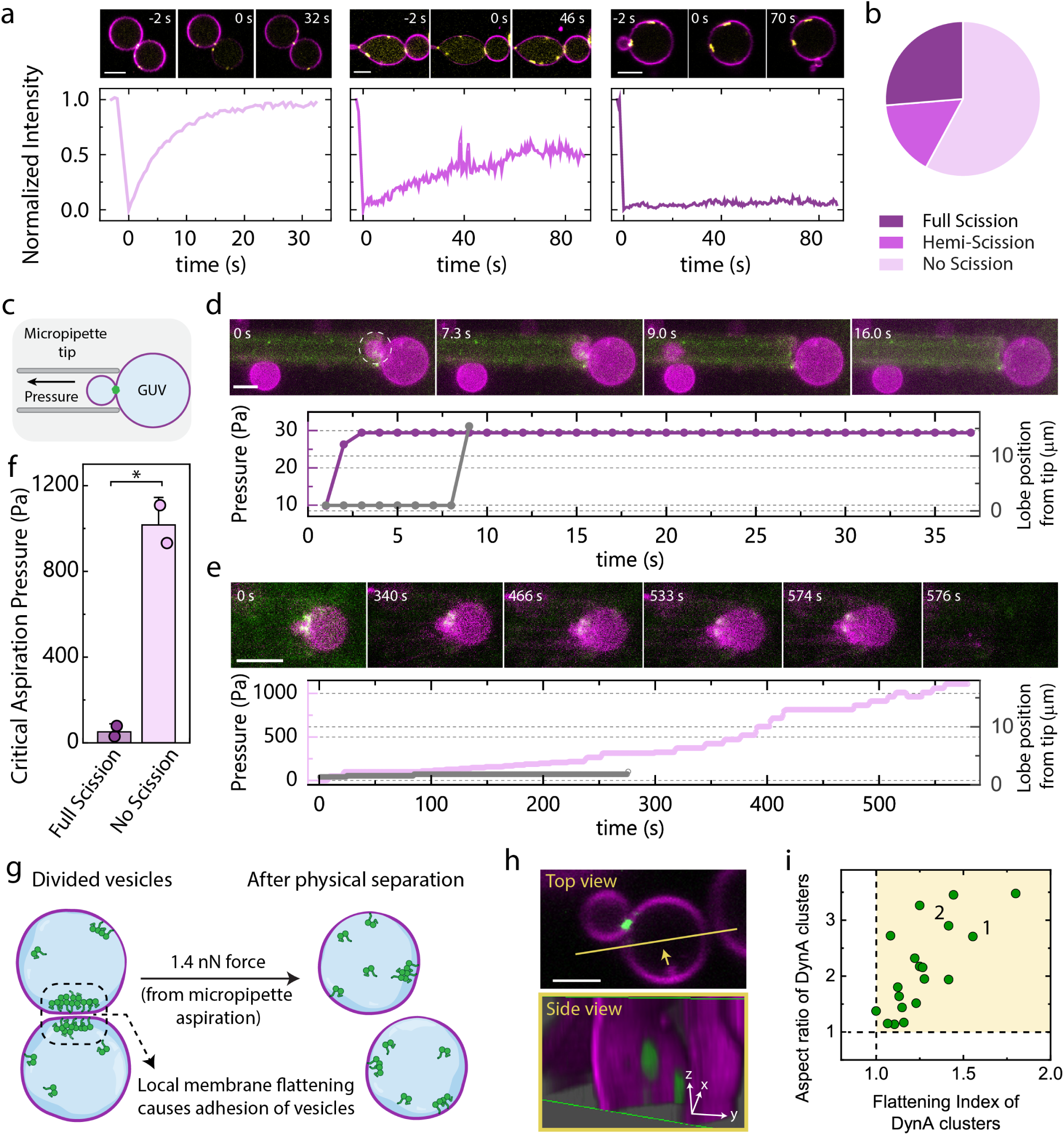
Vesicles divided by DynA adhere to each other via weak physical forces and separate under small, applied force. **a)** Verification of membrane abscission functionality of DynA-mVenus based on FRAP analysis of the membrane connectivity at the necks of dumbbells. Left: full recovery of fluorescent lipids to a normalized intensity Ni ≈ 1 upon photobleaching of one lobe of a chain of dumbbells indicate that no scission has taken place; middle: partial recovery with Ni ≈ 0.5 indicate hemi-scission; right: no recovery with Ni ≈ 0.5 indicate full scission. **b)** Pie chart indicating the fraction of full scission, hemi-scission and no scission events (n = 26 from 3 independent preparations). **c)** Schematic illustration of the micropipette aspiration set-up used to estimate the forces holding the two lobes of a dumbbell together, with one lobe held within the pipette while the other remains outside. The pressure was applied in a controlled stepwise manner over time until the lobes either separated from each other or the entire dumbbell end up getting aspirated and eventually sucked within the tip. The pressure recorded at this time is called critical aspiration pressure (P_C_). The distance between the center of the lobe that is held within the micropipette, and the micropipette tip is used as a guide to track its position over time during the pressure application. **d)** and **e)** Two distinct representative traces of pressure and position versus time reveal a low-pressure regime (d) with P_C_ of 54 Pa and a high-pressure regime with P_C_ of1020 Pa. The pressure and position traces are supplemented with representative fluorescence images over time. Scale bar: 5 µm. **f)** All the recorded P_C_ from low regime (54 ± 35 Pa, n = 2) and high regime (1020 ± 125 Pa, n = 2) are represented as a bar plot. The data is collected from two independent experimental repeats. The statistical significance was tested using two sample t-test after Welch correction (**p < 0.01). **g)** Force of 1.4 nN calculated from the low-pressure regime match weak van der Waals forces, which we hypothesize arises because of DynA-mediated flattening at the post-division interface. **h)** 3D reconstructions of a confocal images reveal an elongated disc-like architecture of DynA (green) enriched at the membrane neck (magenta) of a dumbbell. The top panel shows the top view (xy scan) of the dumbbell, and the bottom panel shows an orthogonal optical section (side view) reconstructed from the same dataset along the yellow line, with the viewing direction indicated by the arrow. Scale bars: 2 µm (top view); 2µm, 1µm, and 0.6 µm for x, y, z dimensions respectively (side view) **i)** Quantitative analysis of 3D DynA clusters (n = 19, N = 2). The three principal dimensions of each cluster, ordered by increasing length (L_min_, L_mid_, and L_max_) were used to estimate the aspect ratio, 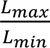 and flattening index, 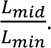 Ratios greater than 1 indicate elongated and flattened cluster geometries, respectively. A total of 19 clusters were analyzed.

Surprisingly, we observed that after full division, the two lobes of the dumbbell almost always remained adhered to each other. To quantify these residual interactions, we measured the adhesive forces between the lobes using a micropipette aspiration (MPA) assay. MPA experiments were performed on dumbbell-shaped vesicles with DynA at the neck, where one lobe of the dumbbell was held within the pipette while the other remained outside (Figure 4c). Pressure was applied and slowly increased until the two lobes separated from each other (or the entire dumbbell was aspirated and pulled into the micropipette tip). The corresponding pressure at this transition point is defined as the critical aspiration pressure (P_c_). Strikingly, we observed two distinct regimes: A low-pressure regime (Figure 4d, Figure S13) with P_c_ = 54 ± 35 Pa (or force = 1.4 ± 0.9 nN; mean ± SD) where the lobes are readily separated from each other; and a high-pressure regime (Figure 4e, Figure S14) that was characterized by a 20-fold larger value of P_c_ = 1.02 ± 0.12 kPa (or force = 24 ± 3 nN). The low forces match typical non-specific binding forces (0.4-1.6 nN) between membranes that were reported before,^53^ such as van der Waals forces, electrostatic attraction or depletion effects. In contrast, the higher forces of order 24 nN are more characteristic of irreversible or specific ligand-receptor binding interactions such as streptavidin-biotin or integrin-fibronectin mediated cell adhesion on a surface.^53,54^

These distinct regimes point to different states of the dumbbell necks. Dumbbells in the high-pressure regime likely retain an open, connected neck indicating that scission has not occurred, whereas those in the low-pressure regime have undergone scission but remain weakly associated through non-specific adhesion (Figure 4f). These weak interactions are unlikely to be purely electrostatic, as vesicle adhesion persisted even after removal of Mg^2+^ ions from the OAS (Figure S15). Instead, we hypothesize that weak interactions can arise because of the DynA mediated flattening of the membranes at the post-division interface increasing the contact area between the two lobes, thereby amplifying short-range attractive interactions such as van der Waals and depletion forces that can stabilize vesicle adhesion even after scission (Figure 4g).^55^ If this interpretation is correct, one would expect to observe flattened, elongated DynA assemblies at the interface of the divided GUVs. We indeed observed, disc-like DynA in 3D confocal images of dumbbells after full division (Figure 4h, Figure S16). We quantitatively assessed the 3D shape of the DynA clusters in terms of the three dimensions defining each cluster (L_min_, L_mid_, and L_max_). DynA clusters exhibited an aspect ratio 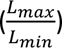 and flattening index 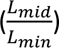 significantly greater than 1, confirming their elongated and flattened, disc-like architecture rather than a spherical cluster (Figure 4i).

### The D1 subunit of Dynamin A suffices for division of synthetic cells

Although DynA is encoded as a single gene, and thus significantly simplifies the division machinery relative to multi-component division machineries found in nature such as the ESCRT-III system, it is nevertheless a large protein (137 kDa) for inclusion in a minimal synthetic genome, which must remain compact to maintain engineerability, stability and biosynthetic capacity.^56,57^ Notably, our DynA contains two bacterial dynamin-like domains at the N-terminal D1 (residues 1-609, 71.2 kDa) and C-terminal D2 (residues 561-1193, 74.4 kDa). In *B. subtilis*, these subunits fused from separate dynamins in its evolutionary past, now jointly cooperating to maintain membrane integrity under various environmental stresses.^58^ Only the D1 subunit was found to associate with the membrane via its membrane binding region, whereas D2 preferably localized within cytoplasm.^26,44^ Indeed, membrane tethering is governed exclusively by the lipid binding domain (residues 360-367) in the D1 subunit.^24,26^ Given this exclusive functionality of D1, we asked whether the D1 subunit alone is sufficient to drive membrane abscission. To test this, we reconstituted purified D1 subunit in pre-deformed GUVs and found that it indeed localized at the necks of dumbbell-shaped GUVs and, most importantly, also induced membrane abscission (Figure S17a, b). By contrast, the D2 subunit did not form any neck enrichments, consistent with the absence of a lipid-binding motif (Figure S17c).

To test if the D1 subunit of DynA can be genomically encoded for synthetic cell abscission, we designed a DNA construct where we fused mVenus at the C-terminal of D1 (Figure 5a). Expressing this D1-mVenus (99.9 kDa) construct in the presence of fluorescently labelled lysine residues yielded, as expected, only a single fluorescent band at ∼100 kDa corresponding to full length D1-mVenus (Figure 5b) but no truncated polypeptides. We also confirmed the correct folding and maturation of mVenus within D1-mVenus during expression in PURE by monitoring its bulk fluorescence emission (Figure 5c). We observed a remarkable 85-fold increase in fluorescence, reaching steady-state within 2.5 hours (about 0.8 hour earlier than for DynA-mVenus expression; Figure 1d, Figure 5c, Figure S3). Expression of D1-mVenus (873 amino acids), with its shorter open reading frame, thus proceeded faster and yielded six-fold higher (Figure S18) protein levels compared to DynA-mVenus (1457 amino acids).^22^ When expressed within GUVs containing 30% DOPG and 100 mM sucrose, we observed a clear time-dependent increase in fluorescence intensity, indicating successful production of D1-mVenus (Figure 5d, f). Coherent with the bulk expression, D1-mVenus reached maximum expression levels 0.4 h earlier than DynA-mVenus expression within the GUVs (Figure S19).

**Figure 5.**
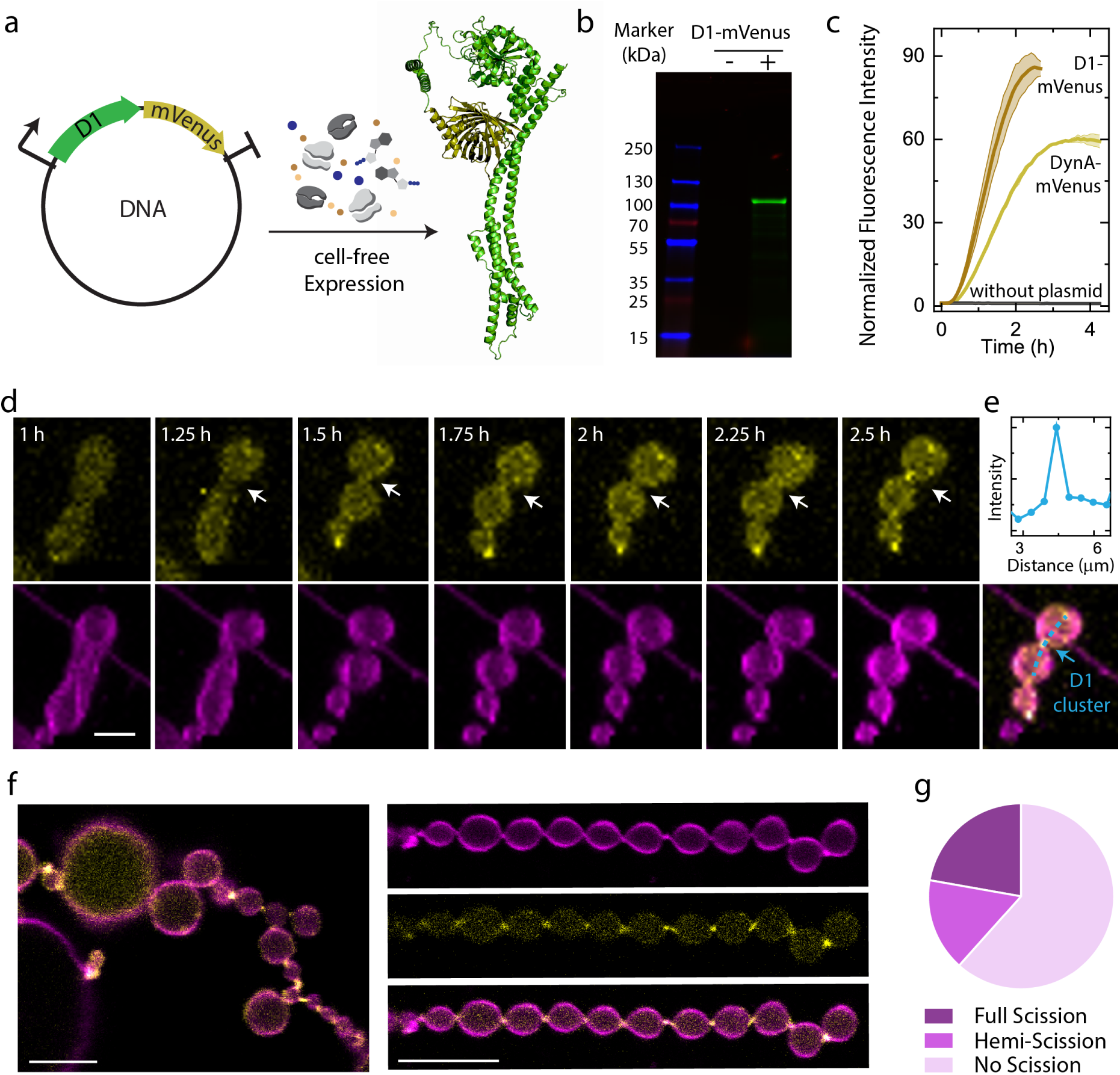
Minimal D1 subunit of DynA is sufficient to program genomically encoded Synthetic Cell division. **a)** Schematic illustration of in vitro expression of D1-mVenus using PURE cell-free expression system. The D1 subunit of DynA is fused with mVenus at the C-terminal causing no structural differences as confirmed with AlphaFold3 predictions. **b)** In vitro expression from D1-mVenus or DynA-mVenus template (both 10 ng/uL) was carried out in the presence of GreenLys and the translated products were analyzed by fluorescence imaging of SDS-PAGE gel. **c)** Time-resolved normalized fluorescence of D1-mVenus during cell-free expression. The fluorescence intensity at time, t is normalized with respect to intensity at t = 0. The results represent an average of two independent measurements; the shaded region depicts the SD. **d)** Time-lapse series showing 3D maximum intensity projections of confocal Z-stacks of a pre-deformed vesicle expressing D1-mVenus, which transforms into chain of dumbbell with D1-mVenus enriched at a newly formed neck (indicated by arrow). **e)** Line-segment analysis shows D1-mVenus enrichment at the two newly formed necks. Scale bar: 2 µm. **f)** Representative image of in vitro expressed D1-mVenus encapsulated within GUVs (large field of view on the left and an example of a chain of dumbbells on the right) prepared using 30% DOPG and 100 mM sucrose. 20 ng/µL of D1-mVenus DNA template was used. Scale bar: 5 µm (left panel) 10 µm (right panel). **g)** Pie chart indicating the fraction of full scission, hemi-scission and no scission events at the necks of dumbbells enriched with D1-mVenus (n = 24, from 2 independent preparations).

Notably, D1-mVenus began enriching at curved membrane regions within 40 minutes of expression and the protein localized to the necks of dumbbells and chains thereof, exhibiting an enrichment ratio of 5.5 (Figure 5d, Figure S11). This confirms that D1 confers the curvature-sensing ability to DynA. To assess whether genomically encoded D1-mVenus can independently drive abscission, we examined the membrane connectivity at dumbbell necks using FRAP (Figure 5g). D1-mVenus induced membrane remodeling events within 38% of the necks, of which 21 % culminated in full scission and 17 % in hemi-scission state. The D1 subunit alone therefore has a comparable membrane remodeling capability to that of full-length DynA. It was previously reported that membrane tethering by D1 can be unstable and has a higher chance of reversibly going back to open neck state as compared to DynA,^24^ but this difference does not appear to intervene with the functionality of the D1 subunit of DynA as a minimal genomically encoded synthetic-cell abscission machinery.

## Discussion and conclusions

Abscission constitutes the final step of division, where the two constricting leading edges of the membrane undergo fusion to pinch a single neck of a dumbbell GUV into two independent, sealed daughter cells. It is the final link in the life cycle of a (synthetic) cell. While abscission was previously explored by reconstituting purified DynA, we here showed for the first time an autonomous abscission machinery based on genome-encoded DynA. Although cell-free protein expression systems such as PURE can pose limits for synthesizing full-length proteins above 100 kDa, we showed that functional DynA-mVenus with a molecular weight of 163 kDa can be synthesized without detectable truncated C-terminal fragments. Cell-free expression of dynamin from its DNA template was found to enrich at the membrane neck of GUV dumbbells and induce scission to complete division.

Membrane peripheral proteins, including DynA, perform diverse biological functions such as structural scaffolding and shaping (e.g., BAR domain proteins), intracellular signaling (e.g., Ras, Rab, Rho subfamilies), cellular defense and membrane repair (e.g., Annexins, DynA). Together, these proteins constitute 5% of the human^59^ and 19% of the bacterial^60^ proteome suggesting the inevitable requirement of this family of proteins for reconstituting similar functions within synthetic cells. It is therefore critical to establish strategies for their successful genomic coding and cell-free expression, especially for DynA, which is prone to aggregation owing to its hydrophobic lipid binding loop. Our results showed that introducing negatively charged lipids such as DOPG facilitated the expression of functionally active DynA. Such findings are directly applicable to other membrane peripheral proteins where negatively charged lipids such as DOPG for bacterial proteins and DOPS for eukaryotic proteins may help preclude aggregation.

Whereas previous reports established dumbbell shapes by integration of cholesterol-labeled DNA nanostructures in the outer leaflet of GUVs,^10^ we here formed dumbbell-shaped GUV using sugar asymmetries in the inner/outer buffers. In line with studies from the Lipowsky group,^34^ our results showed that such asymmetric-sugar-induced GUV deformation generates significant positive spontaneous curvature and produces approximately 40% dumbbell-shaped GUVs. This avoids the unnecessary modification of chemical composition and physical properties of the membrane (such as rigidity and fluidity) from cholesterol and the steric effects introduced by DNA nanostructures, thereby keeping the system minimal and compatible with other synthetic cell modules.

FRAP experiments demonstrated that DynA-enriched necks in dumbbell GUVs culminated in full scission in 27% of cases and hemi-scission in 15%. These outcomes highlight that a genomically encoded, in vitro-expressed DynA can execute membrane remodeling events. Several factors may explain why the remaining DynA-enriched open necks did not undergo membrane remodeling, likely reflecting a complex interplay between DynA activity and membrane mechanics. For example, membrane tension, regulated by the osmotic conditions in the extracellular medium, is known to strongly influence membrane scission, as demonstrated for eukaryotic Dynamin at clathrin-coated pits during endocytosis.^61^ In our system, we found that changing the osmotic pressure difference between OAS and IAS from the previously reported ratio of 1.30,^10^ to 0.92 increased the scission events from 26% to 76% when using purified DynA (Figure S20a). Using a FliptR fluorescent lipid tension probe, we also detected an increased membrane tension under these osmotic conditions (Figure S20b). According to elastic-barrier models, increased membrane tension promotes membrane fusion by bringing opposing bilayers into closer proximity and lowering the energy barrier.^15^ Applying this model to DynA-induced abscission, we suggest that increased tension constricts the dumbbell neck, thereby acting synergistically with DynA to facilitate scission.

It is also plausible that the abscission functionality of DynA is negatively affected under in vitro transcription-translation conditions. When purified DynA was reconstituted in the same cell-free conditions used for genomic encoding, but without DNA, it induced abscission in only 25% of events (instead of 76%; (Figure S20a), very similar to the 27% observed for cell-free-expressed DynA-mVenus (Figure S21). This observation suggests that the cell-free system affects DynA performance which is unsurprising since PURE contains ∼70 individual components (enzymes, nucleotides, amino acids, etc) that could conceivably interact with DynA. We suspect the high levels of GTP (∼3 mM) in cell-free systems to be one of the main reasons. Indeed, when purified DynA was reconstituted with GTP, the abscission activity was strongly reduced (by 69%) compared to GTP-free conditions, but it recovered to nearly maximum scission efficiency (71%) after ∼3 h, likely reflecting the GTP hydrolysis cycle of DynA (Figure S22). Because the PURE energy regeneration module keeps GTP levels high, GTP may reduce scission yields of cell-free-expressed DynA. Interestingly, both D1 and D2 subunits contain GTP-binding domains, but membrane remodeling by purified DynA (both fusion and fission) was found to be GTP independent.^26,44^ Our results suggest that DynA-mediated membrane remodeling may be regulated by local concentrations of GTP, determining when DynA enters its active function. Due to the evolutionary relationship between bacteria and mitochondria and the homology of bacterial dynamins and mitofusins,^26^ these findings can be relevant to the mechanism of mitochondrial fusion.

Our findings revealed that the minimal D1 subunit of DynA, which houses its lipid binding domain, can also be genomically encoded and independently induce membrane scission. Its much smaller gene size (being nearly half the size of full-length DynA) reduces the burden on the synthetic genome,^56,57^ while enabling faster expression and a ∼5-fold higher protein yield. These features make D1 well-suited for integration into more complex synthetic cell designs. Furthermore, the D1 subunit can potentially be even further minimized in future work, for example by removing the GTPase domain, which may not be required for membrane scission at all.

A central challenge in the bottom-up synthetic cell field is sustaining division of synthetic cells over multiple cycles. Now that the synthetic cell abscission has been genomically encoded, it can be coupled to a genetic oscillator to generate periodic bursts of DynA expression, enabling the timely assembly and disassembly of the abscission machinery during each division cycle.^62^ As daughter vesicles are smaller after DynA-induced division, with less excess membrane and higher curvature, subsequent divisions will be increasingly difficult, and division must therefore be coupled to growth, for example through phospholipid synthesis^5^ or vesicle fusion.^4^ Currently, division relies on externally imposed osmotic changes to generate dumbbells. An alternate and more active approach can involve shape-generating machinery. While FtsZ can assemble into rings and constrict GUVs into dumbbell geometries,^63^ these rings typically stall without progressing to sub-100nm sizes and division^63^ – emphasizing the need for an abscission machinery.

Overall, we have shown that division of a dumbbell shaped vesicle can now be programmed autonomously by using just a one-component DynA machinery. The simplicity of DynA and particularly its minimal subunit D1 enables facile integration with other synthetic cell modules such as DNA replication, constriction, and metabolism without imposing a high load on the synthetic genome – establishing an important step towards making a synthetic cell that is able to sustainably replicate itself.

## Data availability

The raw data and processed data for reproducing the figures are deposited in Zenodo (10.5281/zenodo.21453617) and will be made publicly available upon publication of the manuscript.

## Code availability

The image analysis source code is available via Zenodo (10.5281/zenodo.21453617) and will be made publicly available upon publication of the manuscript.

## Acknowledgment

We thank Marc Bramkamp for kindly providing the plasmid for DynA; Liedewij Laan for kindly providing tRNA mix and amino acids; Eli van der Sluis and Ashmiani van den Berg for purification of DynA and D1; Jeffrey den Haan and Sonam Marapin for technical support.; and Federico Ramirez Gomez, Marcos Arribas Perez, Bert van Herck, and Rafael B. Lira for helpful discussions and advice regarding experiments. We gratefully acknowledge financial support from The Netherlands Organization of Scientific Research (NWO/OCW) Gravitation Program Building a Synthetic Cell (BaSyC).

