## Supplementary material for "Division of Synthetic Cells Using a Genomically Encoded One-Protein Divisome": Methods and Supplementary Figures: Genomically encoded Synthetic Cell Abscission_SI.pdf

### Reagents

All reagents were ordered from Sigma Aldrich unless stated otherwise. DOPC (1,2-dioleoyl-sn-glycero-3-phosphocholine) (850375), DOPE-PEG(2000) amine (1,2-dioleoyl-sn-glycero-3phosphoethanolamine-N-[amino(polyethylene glycol)-2000] (ammonium salt)) (880234), 18:1 (Δ9-Cis) PG (1,2-dioleoyl-sn-glycero-3phospho-(1'-rac-glycerol) (sodium salt)) (840475) and DOPE-rhodamine (1,2-dioleoyl-sn-glycero-3-phosphoethanolamine-N-(lissamine rhodamine B sulfonyl) (ammonium salt)) (810150C), 18:1 Cyanine 5 PE (1,2-dioleoyl-sn-glycero-3-phosphoethanolamine-N-(Cyanine 5)) (810335) were purchased from Avanti Lipids. Lipids were resuspended in anhydrous chloroform (288306, Sigma-Aldrich) and stored under argon at -20 °C. Silicone oil (viscosity 5 cst (25 °C), Carl Roth, 7844.1), mineral oil (BioReagent, M5904), n-decane (99+%, pure, Acros Organics), chloroform (≥99% pure, Sigma Aldrich, 1024470500), sucrose (Sigma Aldrich, S0389), glucose (G7021), $MgCl_2$ (M8266), OptiPrep (60% (w/v) iodixanol in water; D1556) were required for GUV preparation. Bovine Serum Albumin and β-casein used for the passivation of glass coverslips were purchased from Thermo Fisher. PURE*frex*2.0, DnaK mix, and GroE mix (GeneFrontier Corporation, Chiba) were stored and handled according to the manufacturer's instructions. SUPERase·In™ RNase Inhibitor (AM2696) for inhibiting RNases in PURE reaction mix was ordered from Thermo Fisher. Flipper-TR for fluorescence lifetime imaging was ordered from Spirochrome AG (SC020).

### Protein Purification and labelling

Purification and labelling of DynA, D1, and D2 was performed as described previously.[1]

### Preparation of DNA constructs

pET16b-DynA (9.2 kbp, provided by Bramkamp lab[2]) and pED101-D1 (7.5 kbp, provided by Bramkamp lab[2]) plasmids were linearized by standard PCR with KOD Xtreme Hot Start DNA Polymerase (71975-3) using primers FW-DynA and RV-DynA (Table 4). The plasmids already contained a T7 promoter, terminator and ampicillin resistance gene. mVenus gene fragment was linearized by standard PCR with Phusion High Fidelity DNA polymerase (Thermo Fisher Scientific) from pWKD014 DNA with primers FW-mVenus and RV-mVenus. Both sets of primers contain overhangs for further assembly of DynA or D1 with mVenus via Gibson Assembly. The PCR products were incubated with DpnI (New England BioLabs®, Inc.) to remove residual plasmid, as confirmed by testing on a 0.7% agarose gel stained with SYBR safe, imaged with a ChemiDocTM Imaging System (BioRad Laboratories). The products were purified with the innuPREP PCRpure Kit (Innuscreen, 845-KS-5010250). DNA concentration and purity were measured using a ND-1000 UV-Vis Spectrophotometer (Nanodrop Technologies). Gibson assembly (Gibson Assembly® Master Mix of New England BioLabs®, Inc.) was performed with 3-fold molar excess of mVenus fragment with respect to DynA or D1 fragments for 1 h at 50 °C. Transformation of the Gibson assembly products into E. coli competent cells (NEB® 5-alpha Competent E. coli (High Efficiency)) was done by heat shock, after which cells were resuspended in 950 µL of fresh prechilled liquid lysogeny broth (LB) medium and incubated for 1 h at 37 °C and 250 rpm. Then, the cultures were plated in solid LB medium with ampicillin and grew overnight at 37 °C. Colonies were picked up and cultured in 5 mL of liquid LB medium with 100 µg/mL of ampicillin for 16 h at 37 °C and 250 rpm. Plasmid purification was performed using the PureYield™ Plasmid Miniprep System (Promega). Concentration and purity of DNA were checked on NanoDrop.

### Cell-free expression and gel/fluorescence analysis

PUREfrex2.0 reaction mixture was assembled in a PCR tube by combining all the components listed in Table 2. The reaction mixture was supplemented with 1 µL of GreenLys reagent (FluoroTectTM GreenLys, Promega) and cell-free expression was performed at 37 °C for 4 h. Lysine residues constitute 9% of amino acid content of DynA-mVenus and are distributed throughout the structure, making it well suited for GreenLys labeling. Sample was treated with 0.2 µL of RNase (RNaseA Solution, 4 mg/mL, Promega, A7973) for 30 min and proteins denatured for 5 min at 95 °C in 1× SDS loading buffer (non-reducing, 4X, Thermo scientific, J63615.AC) with 10 mM dithiothreitol (DTT). Samples were loaded on a 4-15% SDS-PAGE gel (BIO-RAD, 4568085). Visualization of the fluorescently labeled protein was performed on a fluorescence gel imager (Typhoon, Amersham Biosciences) using 488, 532, and 633 nm lasers. The gels were subsequently stained with InstantBlue® Coomassie Protein Stain (abcam, ab119211) for 10 min and washed with Milli-Q water for 3 h, with the water exchanged every hour to remove residual stain. Gels were subsequently imaged with a ChemiDocTM Imaging System (BioRad Laboratories) and the images were analyzed using ImageJ Gel Analysis tool.[3]

For cell-free expression experiments in GUVs, a Home-made PURE Solution I was used as outer aqueous solution. Based on the previously described composition of Solution I,[4] Home-made PURE Solution I was prepared as follows:

An 18-amino-acid mixture was first prepared, containing all standard amino acids except cysteine and glutamine. Because of their different solubilities in water, each amino acid was dissolved in a different concentration of KOH solution to prepare 1000 mM stock solutions. For example, all amino acids except Alanine, Arginine, Asparagine, Aspartic acid, Tryptophan, and Tyrosine were dissolved in 1000 mM KOH solution at a concentration of 1000 mM. Alanine and Arginine were dissolved in 5 mM KOH whereas Asparagine, Aspartic acid, Tryptophan, and Tyrosine were dissolved in 5000 mM KOH. The 18 amino acids were then mixed to obtain a final concentration of 50 mM for each amino acid. Similarly, 50 mM cysteine and glutamine solutions were prepared separately by diluting the respective 1000 mM stock solutions prepared in 5000 mM KOH. 18-amino acid mixture and individual solutions of cysteine and glutamine were aliquoted and stored at -80 °C until further use (stable for upto two years, based on our observations).

Next, all components required for Home-made Solution I were combined in the order listed (Table 1). First, HEPES-KOH, potassium glutamate, magnesium acetate, and spermidine were mixed together with the required volume of MilliQ water to compensate for the remaining volume. A total of 0.2% acetic acid (v/v) in Home-made Solution I was required to adjust the final pH of the buffer to 7.6. Half of this amount was added at this step. Freshly prepared creatine phosphate, folinic acid calcium salt hydrate, dithiothreitol, and all NTPs were then added, followed by the remaining half of the acetic acid solution. Finally, freshly thawed 18 amino acid mix, cysteine, glutamine solutions and tRNA mix were added, followed by addition of glycerol. The pH of the final solution was confirmed with a pH indicator strip (VWR, 662-0200, pH 2.0-9.0) to be ~7.5. The Home-made Solution I was aliquoted and stored at -80 °C for further use and was consumed within 3 months of preparation.

### Lipid-in-oil suspension

Lipid-in-oil suspensions were prepared as described in previous work from Koenderink lab.[5] Briefly, the lipids, DOPC, DOPE-PEG(2000) amine, DOPG, and 18:1 Cyanine 5 PE (or Rhodamine DOPE) were mixed in a desired molar ratio in a 20 mL glass screw neck vial. The amount of each lipid was calculated such that, upon dissolution in the oil mixture, the final total lipid concentration was 0.2 mg/mL. After desiccation using a gentle nitrogen flow, the vial was brought inside a glovebox to prepare a lipid-in-oil suspension under controlled humidity conditions that prevent premature lipid hydration. Inside the glove box, the lipid film was resuspended in 25 µl of chloroform and 400 µl of n-decane. A mixture containing 5.7 mL silicone oil and 1.3 mL mineral oil was then added dropwise to the lipids while vortexing. After tightly closing the vial, the lipid-in-oil suspension was vortexed during an additional 2 min. The lipid-in-oil suspension was removed from the glovebox, tightly capped, sealed with Parafilm, and placed in a beaker containing ice which was sonicated for 15 min while keeping the bath temperature below 30 °C. The mixtures were used the same day in experiments.

### Dumbbell GUV preparation

GUV dumbbells were prepared using a slightly modified inverse-emulsion method.[1] For preparing dumbbells encapsulating DNA templates and the PURE system, Inner and Outer Aqueous Solutions according to Table 2 were prepared. First, 300 µL of lipid-in-oil suspension was gently added to 20 µL of Inner Aqueous Solution (IAS) and mixed by pipetting up and down 30-40 times (without introducing air-water interfaces) to form water-in-oil droplets with lipid monolayer. This emulsion was immediately and carefully layered onto 44 µl of outer aqueous solution (OAS) placed in a 1.5 mL centrifuge tube kept on ice. Upon centrifugation at 400 rcf speed for 15 min at 4 °C, droplets traversed the oil-water interface, acquiring a second lipid leaflet thereby forming GUVs. The top layer was discarded and the bottom layer containing the GUVs were collected and incubated at 37 °C in microscopy chambers (Ibidi, 81817) passivated with 10 mg/mL of BSA in water. Dumbbells encapsulating recombinant DynA or D1 were prepared following the same procedure, except using IAS and OAS compositions as described in Table 3.

### Bulk fluorescence measurements

The temperature-controlled fluorescence measurements were performed on a BioSPX Synergy H1 microplate reader using Greiner plates (384w, F, µClear®, 10-130µl/w, 11mm²/w, polystyrene, black, per 10 pieces, binding: MEDIUM). Excitation and emission wavelengths for mVenus were 500 and 539 nm, respectively.

### Confocal Microscopy

Fluorescence images were acquired on an inverted Leica Stellaris 8 FALCON laser scanning confocal microscope using LasX software (version 4.8.2). During imaging, a constant temperature was ensured by using a box incubator (Okolab). Specifically, a temperature of 37 °C was maintained during cell-free expression of DynA-mVenus or D1-mVenus, whereas a temperature of 25 °C was used for experiments with recombinant proteins. Samples were illuminated using a 63x glycerol immersion objective (HC PL APO CS2 63X/1.30, Leica). Alexa 488, mVenus, Rhodamine, Cy5 were excited at wavelengths of 488 nm, 515 nm, 579 nm, and 649 nm respectively using a white light laser (WLL) at a repetition rate of 80 MHz. The emission light was passed through a 104 µm pinhole and collected in the spectral range of 496 nm-533 nm for Alexa 488 and a range of 520 nm-568 nm for mVenus using a HyDX1 detector operated in digital mode with an internal gain of 65 respectively. Emission light for Rhodamine was collected in the spectral range of 586 nm-733 nm and for Cy5 in the range of 658 nm-811 nm using a HyDS2 detector operated in analog mode.

***Fluorescence recovery after photobleaching experiments***
For FRAP experiments, both the protein channel and membrane channel were acquired, whereas only the membrane channel was photobleached. Rhodamine-DOPE and Cy5-DOPE were bleached using same settings except using different laser wavelengths. The white light laser was operated under similar settings as described above with a pixel dwell time of 3.53 µs during acquisition of the pre- and post-bleach images. Bleaching was performed on a circular region of interest (ROI), which encompassed one of the lobes of a dumbbell connected via DynA/D1 enrichment (Figure 21, supplementary information). ROIs were thus different in size and aspect ratio for each lobe of dumbbell. The ROI was bleached using two laser lines for Rhodamine-DOPE (571 and 579 nm) or for Cy5-DOPE (639 and 649 nm) operated at 100% intensity each, to efficiently bleach the membrane within three frames (= 875 ms). We acquired three images before bleaching the GUV at four frames per second (4 fps). After bleaching, we acquired 75 frames (per channel) at 4.6 fps to capture membrane dynamics for 32.4 s. The FRAP function of the Leica LAS X software was used to quantify fluorescence recovery in FRAP experiments. The software enabled selecting regions of interest (ROIs) corresponding to the bleached lobe and control lobe and tracking the average intensity over time of each region. The resulting data were then exported, plotted and analyzed. For purposes other than fluorescence recovery analysis, images were processed using ImageJ (v. 1.5.4j). Due to incorrect parsing of .csv files after exporting them from the LAS X software, a custom python script was written to fix their formatting. Another script was written to quantify normalized fluorescence recovery and classify results of FRAP experiments in bulk into full scission, hemi-scission, and no scission state, given an output from the first script. The script output was manually reviewed together with the corresponding FRAP image sequences to verify the classifications and exclude outliers. The normalized fluorescence intensity is defined in the following way for analysis:

With $F_{raw\ data}$ the intensity of the bleached lobe and $F_{raw\ control}$ the intensity of the control lobe,

$$F = \frac{f - f_{post-bleach}}{f_{steady\ state} - f_{post-bleach}}, with\ f = \frac{F_{raw\ data}}{F_{raw\ control}}$$

Normalized curves were obtained as follows: $N = F.\frac{1}{F_{pre-bleach}}$.

Finally, for model fitting, the least squares method using the Levenberg-Marquardt algorithm was used.[6]

Criteria for excluding data points in FRAP analysis during image acquisition were (1) the presence of lipid clusters at the neck, (2) convoluted chains of dumbbells that precluded the visual determination of connectivity between lobes, (3) lobes smaller than ~2 µm that precluded accurate bleaching, and (4) lobes moving up and down in the z direction precluding accurate bleaching.

#### *Fluorescence lifetime imaging microscopy*

For FLIM measurements of membrane tension via the Flipper-TR reporter,[7–9] samples were excited at 488 nm using 2% laser power. Emission was collected in the 575–625 nm range using a HyD detector in photon-counting mode. The laser power was adjusted to ensure a photon count rate below 0.5 photons per excitation pulse to minimise pile-up artifacts, where more than 1 photon per pulse are detected. All acquisitions were performed at a laser repetition rate of 20 MHz. Fluorescence decay curves were fitted using a double-exponential reconvolution model, employing the instrument response function (IRF) generated by the FALCON FLIM module within a fitting window of 0.2–45 ns. In this method, the theoretical decay was convolved with the IRF and iteratively optimized to minimize the difference between the calculated and measured fluorescence decays. Reported lifetime values correspond to the mean intensity-weighted lifetime ($\tau_{m\,int}$), calculated by the software according to the following equation:[9]

$$\tau_{m\,int} = \frac{\sum_{k=0}^{n-1} I[k]\tau[k]}{I_{sum}}$$

where $n$ is set to 2, $I[k]$ is the intensity of each exponential component, $\tau[k]$ the corresponding lifetime, and $I_{sum}$ the total fluorescence intensity.

### Image Analysis

All the representative confocal images, 3D reconstructions, and time-series were exported directly from LasX software (version 4.8.2) post smooth rendering function, which applies interpolation between adjacent pixels to reduce the appearance of jagged edges. Further quantitative image analysis was performed on the corresponding raw confocal images, as described in the following sections:

#### *Classification of DynA-mVenus phenotypes*

Overlay elements of individual GUV were manually extracted from multiple confocal z-stacks for each cell-free expression condition using ImageJ and saved as .csv files. A custom-written python script was then used to crop out all such individual GUVs as tiff files provided as two separate channels (Cy5-DOPE and DynA-mVenus). An ImageJ macro script was written to merge and create a stack of two channels. The cropped GUVs from each condition were manually classified into seven phenotype categories (Figure 2b, main text). Criteria for assigning phenotypes were (1) GUVs without any fluorescent signal (~5-10%) were excluded from the dataset, (2) protein aggregates overlapping with lipid clusters were not regarded as presence of non-ideal or mixed phenotypes but included within the lumen category of functional phenotypes, and (3) the presence of DynA clusters in stomatocyte type necks (membrane topology found in intracellular organelles including nuclear envelope and open autophagosome)[10] were excluded from the dataset.

#### *Quantification of membrane curvatures*

Membrane point curvatures at the tip of the saddle-shaped necks in Figure 3d (main text) were assessed using the ImageJ plugin Kappa - Curvature analysis (version 2.0.0).[11] For each time point of the series, we drew a ~3 µm-wide ROI along the entire saddle shape and used Kappa to extract the membrane curvature in the x-y plane in this region, based on B-splines fit to the membrane contour. The full curves were exported as .csv files and the local curvature strictly at the tip of saddle-shape was used for further analysis as point curvature.

#### *Area Analysis*

Aggregate area in 2D-confocal slices was used as a proxy for the extent of dynamin aggregation across different cell-free expression conditions. A custom ImageJ Macro was written to measure the size of aggregates on a z slice passing through the equatorial plane of majority of GUVs. A fixed threshold of (10-255) was applied on 8-bit images to create binary masks which were further used to analyze the size (area) of the masked aggregates using a size filter of 0.05-5000 $\mu m^2$. Output from different images per condition were merged using another custom-written Macro in ImageJ.

#### *Shape Analysis of DynA clusters*

3D DynA clusters were segmented and analyzed using the 3D Object Counter Plugin in ImageJ.[12] Confocal z-stacks in a single channel (Alexa 488 labelled DynA) were first calibrated using the measured pixel dimensions (width = height = 0.095 µm) and voxel depth (0.33 µm) to ensure accurate voxel dimensions. 3D objects were segmented by intensity thresholding to a fixed value of 10 and analyzed as binary volumes. For each object, the plugin calculated a three-dimensional bounding box defined by the minimum and maximum coordinates occupied by the object along the x, y, and z axes. Width, height, and depth were calculated as the differences between the maximum and minimum object coordinates along the x, y, and z axes, respectively. All measurements were reported in calibrated physical units. The three dimensions were arranged in the order of increasing lengths ($L_{min}$, $L_{mid}$, and $L_{max}$), where ratios, $\frac{L_{max}}{L_{min}}$ and $\frac{L_{mid}}{L_{min}}$ were used to estimate the aspect ratio and flattening index of the DynA clusters, respectively. Ratios larger than 1 mean the clusters are elongated and flattened, respectively.

#### *Estimating number of DynA molecules in a cluster enriched at the neck*

For quantitative imaging, we acquired two channel z-stacks of DynA clusters located at the necks of dumbbells in photon counting mode. As described in the previous section, the 3D DynA clusters were segmented and analyzed using the 3D Object Counter Plugin in ImageJ. For each object, the plugin calculated the volume and mean fluorescence intensity. The concentration of DynA within 3D clusters was extracted using the concentration-intensity calibration curve, and the number of DynA molecules was further estimated using the volume obtained from the plugin.

#### *Estimation of Enrichment ratio*

Fluorescence intensity line profiles were extracted across the dumbbell neck in the LasX software and exported as CSV files. Due to incorrect parsing of .csv files after exporting them from the LAS X software, a custom python script was written to fix their formatting. Line profiles were analyzed using a custom Python script. Intensity profiles were first smoothed using a centered rolling mean (3-point window) to reduce pixel-level noise while preserving the overall peak shape. Each profile was fitted with a one-dimensional Gaussian Amplitude function with a constant background:

$$I(x) = A\, e^{\frac{-(x-\mu)^2}{2\sigma^2}} + B$$

where $A$ is the Gaussian amplitude, $\mu$ is the peak position, $\sigma$ is the standard deviation (cluster width), and $B$ is the constant background intensity. Nonlinear least-squares fitting was performed using the Levenberg-Marquardt algorithm.

The Gaussian Amplitude was used as a measure of protein enrichment at the membrane neck, while the fitted constant background represented the fluorescence intensity within the lumen. The enrichment ratio providing a normalized measure of protein enrichment at the neck relative to the luminal fluorescence was calculated using:

$$Enrichment\ ratio\ (E_R) = \frac{A}{B}$$

DynA-mVenus enriched necks where the enrichment ratio fell below its respective control, i.e., where enrichment was not qualitatively observed, were not considered protein-enriched necks and were therefore excluded from further analysis.

### Micropipette Aspiration Setup

#### *Micropipette Fabrication*

Micropipettes (tip diameter 5–9 µm) were pulled from borosilicate capillaries (ID 0.58 mm, OD 1.0 mm, length 100 mm; Harvard Apparatus) in a two-step procedure. Capillaries were first pulled on a laser-based puller (Sutter P-2000; settings: Heat 450, Filament 4, Velocity 50, Delay 255, Pull 150) to produce closed tips with extended shanks. Tips were then opened and shaped on a microforge (Narishige MF-900) fitted with a 35× air objective, using a U-shaped platinum filament (150 µm) coated with low-melting-point glass (VPS, ~600 °C). Each tip was briefly contacted with the molten glass bead, then retracted after cooling, creating a small opening. Re-immersing the tip, controlled capillary-driven glass flow and set the final diameter, resulting in smooth, perpendicular openings.

#### *Instrumentation and Pressure Control*

Experiments were performed on a Nikon Eclipse Ti microscope with a Crest X-light confocal spinning disk, a Hamamatsu Orca-Flash 4.0 camera, and a Spectra X LED illumination system (Lumencor). A custom aspiration setup was mounted on the microscope stage.[13] Micropipettes were secured in a Narishige holder attached to a micromanipulator for three-dimensional (x, y, z) positioning and adjustment of the insertion angle. Suction pressure was set hydrostatically via a vertically translatable water reservoir (15 mL syringe without plunger) mounted on a motorized translation stage (LTS300, Thorlabs; 30 cm range, 1 µm precision), connected to the holder with C-Flex tubing (ID 2.4 mm, OD 4 mm). Pressure adjustments were performed manually via stage control with custom C-sharp software (developed by Brahim Ait Said from the AMOLF research institute, Amsterdam),[13] and image acquisition was synchronized using NIS-Elements AR 5.02.03 64-bit software.

#### *Aspiration Chamber Preparation*

Glass coverslips (No. 1.5H, 9 × 50 mm, Thorlabs) were cleaned with absolute ethanol and dried under a stream of $N_2$ gas. Aspiration chambers were then assembled using a custom 3 mm-thick aluminum spacer with internal dimensions of 40 × 12 × 1.5 mm (L × W × H).[13] The spacer was sandwiched between two cleaned coverslips fixed with vacuum grease, leaving both long edges open for pipette insertion.

***System Passivation***
Aspiration chamber and micropipette surfaces were passivated with a β-casein solution (5 mg/mL in OAS) to prevent membrane adhesion. A 70 µL droplet was placed at the chamber center to contact both top and bottom surfaces. For the pipette, air bubbles were first cleared from the tubing and holder before backfilling with the same β-casein solution until a droplet formed at the tip. The pipette was subsequently inserted into the chamber at ~4° angle under slight positive pressure to prevent air entry, and flow and tip integrity verified. The assembled system was passivated for at least 30 minutes, then washed twice with OAS buffer.

***GUV Preparation and Loading***
The chamber was loaded with 54 µL OAS and 6 µL GUV suspension was left to sediment for 5 minutes. The chamber was subsequently sealed with ~200 µL heavy mineral oil to prevent evaporation and maintain constant hydrostatic pressure.

***Aspiration Protocol***
No FRAP was performed to predetermine the dumbbells for aspiration. The only selection criteria was for the dumbbells to have two distinct lobes with a DynA cluster at the neck and at least one lobe measuring ~10 µm in diameter. The zero-pressure reference was set by adjusting reservoir height until no tracer flow was observed in the pipette (~10 µm $H_2O$ precision). Pressure was applied in steps by stepwise lowering the reservoir from -0.1 to -250 mm relative to this reference, corresponding to an applied pressure range of 0.1–1.9 kPa. Each step included a 5-second equilibration period. Fluorescence images (100 ms exposure, 60× water-immersion objective, NA 1.0, WD 2.0 mm, pixel size 108 nm) were collected at each step and GUV deformation was monitored throughout. Neck scission was defined as complete loss of membrane continuity between the two lobes. After the pressure ramp, GUVs were released by reversing the reservoir height. Each GUV required approximately 20 minutes to complete.

***Force Estimation***

The apparent force required to separate the two lobes of a dumbbell-shaped GUV was estimated using geometric relation,

$$F = P_c \, A_{pipette},$$

where $P_c$ is the critical aspiration pressures achieved from the pressure versus time traces and $A_{pipette}$ is the cross-sectional area of the pipette tip.[14] This value is reported as an apparent force that establishes an operational upper-bound scale for the system, since it assumes that the pressure acts uniformly across a perfectly flat cross-sectional area and treats the GUV as a rigid object rather than a fluid membrane system. Plugging the micropipette tip radius of 2.87 µm into the equation along with measured values of $P_c$ provides average force levels of 1.4 nN and 24 nN in the low-pressure regime (scission state) and high-pressure regime (open neck), respectively.

### Statistics and Reproducibility

Statistical analysis of the data was performed as indicated in the figure captions. No statistical method was used to predetermine the sample size. All statistical analyses and curve fittings were performed using OriginPro (OriginLab, Northampton, MA, USA), unless otherwise stated.

### Protein structures

DynA-mVenus and D1-mVenus structural predictions were made using AlphaFold3. Further processing was done using PyMOL™ (version 3.0.3). This involved setting the background white, changing the colors of different domains, ray tracing the images by setting the mode to 3 to make them higher quality. The images were further saved by ray tracing at 2400 X 2400 pixels at 300 dpi. To assess the impact of introducing mVenus on Dyn or D1 structure, the root mean square deviation (RMSD) between corresponding atoms of the two superimposed structures was calculated using PyMOL™. An RMSD of 0 Å indicates identical structures, while values below ~1 Å generally reflect very high structural similarity within experimental noise. RMSD values above approximately 2–3 Å are typically indicative of substantial structural differences or conformational changes.[15]

**Table 1.** Composition of Home-made Solution I.

| Component | Supplier | 2X preparation |
|---|---|---|
| **HEPES-KOH (pH = 7.6)** | Sigma Aldrich, H3375 | 100 mM |
| **Potassium glutamate** | Sigma Aldrich, G1501 | 200 mM |
| **Magnesium Acetate** | Sigma Aldrich, M5661 | 26 mM |
| **Spermidine** | Sigma Aldrich, 85558 | 4 mM |
| **Dithiothreitol** | Sigma Aldrich, 43815 | 2 mM |
| **18 amino acid mixture (excluding cysteine and glutamine)** | Provided by L. Laan lab | 0.6 mM |
| **L-Cysteine** | Provided by L. Laan lab | 0.6 mM |
| **L-Glutamine** | Provided by L. Laan lab | 0.6 mM |
| **Creatine Phosphate** | Sigma Aldrich, 27920 | 40 mM |
| **Folinic acid calcium salt hydrate** | Sigma Aldrich, 47612 | 46 µg/mL |
| **ATP, GTP, CTP, UTP** | New England Biolabs, N0450S | 4 mM |
| **tRNA mix** | Provided by L. Laan lab | 112 OD260/mL (4.5 mg/mL) |
| **Glycerol** | Sigma Aldrich, G5516 | 2.5% (v/v) |

**Table 2.** Compositions of inner and outer aqueous solutions used in preparation of vesicle dumbbells encapsulating DNA templates and the PURE system.

| | **Inner Aqueous Solution (IAS),** *total volume = 20 µL* | **Outer Aqueous Solution (OAS),** *total volume = 44 µL* |
|---|---|---|
| **Solution I of PURE kit** | 10 µL | - |
| **Solution II of PURE kit** | 1 µL | - |
| **Solution III of PURE kit** | 2 µL | - |
| **RNase inhibitor** | 0.5 U/µL | - |
| **Sucrose** | 50 – 200 mM (as described in Figure captions) | - |
| **DNA template** | 4-20 ng/µL (as described in Figure captions) | - |
| **Home-made Solution I** | - | 22 µL |
| **Glucose** | - | Same as sucrose concentration used in IAS |

**Table 3.** Compositions of inner and outer aqueous solutions used in preparation of vesicle dumbbells encapsulating purified proteins.

| | Inner Aqueous Solution (IAS) | Outer Aqueous Solution (OAS) |
|---|---|---|
| **Tris-HCl, pH = 7.5** | 50 mM | 50 mM |
| **$MgCl_2$** | 5 mM | - |
| **Optiprep** | 37% (volume/volume) | - |
| **Recombinant DynA or D1 subunit** | 300 nM | - |
| **Glucose** | - | To adjust desired osmolality |

**Table 4.** List of primers used in this study.

| Name | Sequence (5'→3') |
|---|---|
| **FW-DynA** | ATGGACGAGCTGTACAAGTAACACCGCTGAGCAATAACTAGCAT |
| **RV-DynA** | ATATCAAGCTTATCGATACCGTCGACCTCGAAGTGATGGTGATGGTGATGGCT |
| **FW-mVenus** | GGTATCGATAAGCTTGATATCGAATTCCTGCAGATGGTGAGCAAGGGCGAGG |
| **RV-mVenus** | CTAGTTATTGCTCAGCGGTGTTACTTGTACAGCTCGTCCAT |

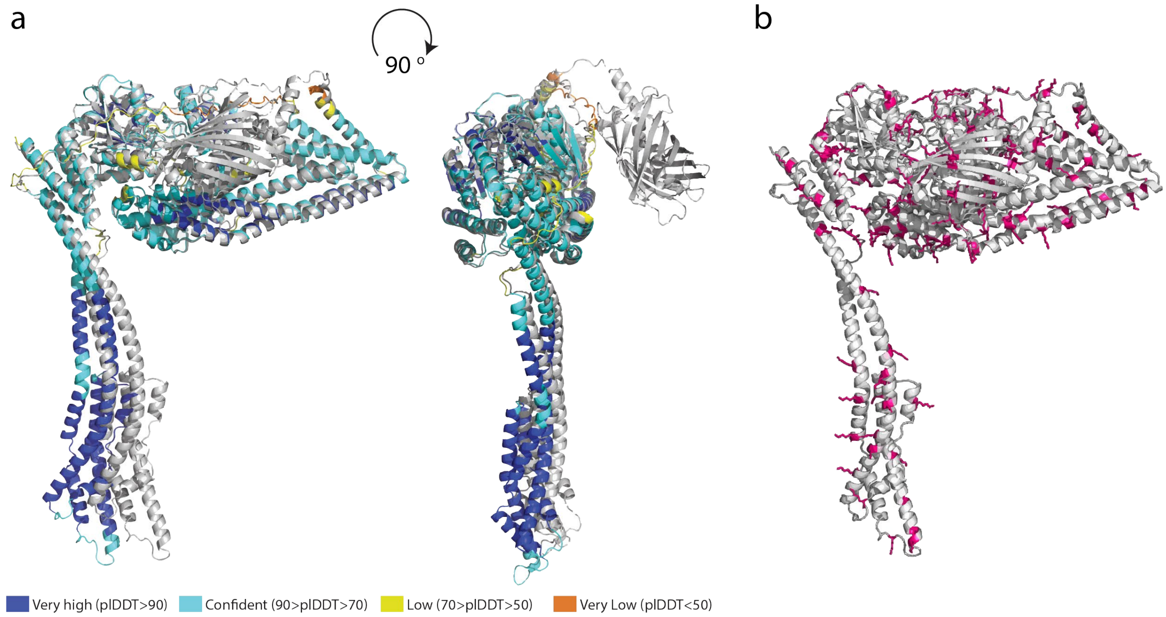


**Fig. S1**. AlphaFold3 structural predictions. a) DynA is colored according to pLDDT score and DynA-mVenus in grey. DynA prediction had a pTM score of 0.75 and DynA-mVenus of 0.65, the pTM score provides an estimate of the global structure accuracy, ranging from 0 to 1. A value of 1 would mean a perfect prediction. pLDDT is a per-atom confidence estimate on a 0-100 scale, where a higher value indicates higher confidence. DynA and DynA-mVenus structural predictions are aligned in PyMOL to reveal an RMSD (root mean square deviation) value of 0.948 Å (<2 Å), indicating no significant structural difference is introduced upon fusing mVenus at the C terminal of DynA. b) Distribution of lysine residues (pink) across DynA-mVenus (grey) structure. Lysine constitutes 9% of amino acid content in DynA-mVenus.

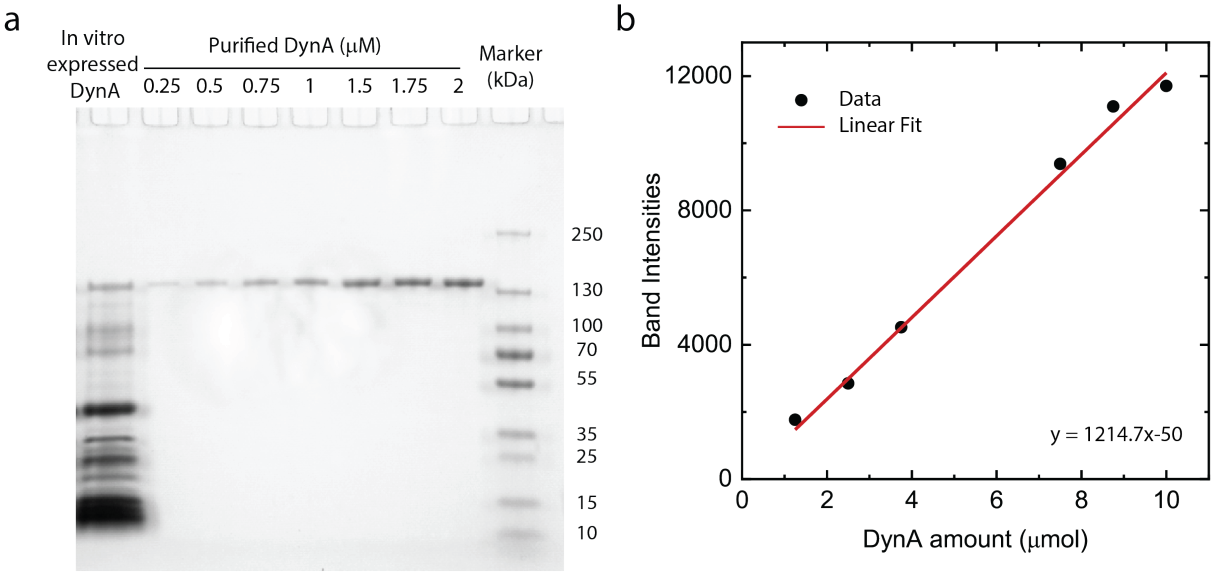


**Fig. S2.** Measurement of cell-free expressed DynA concentration using the PURE system. a) SDS-PAGE analysis of translated products obtained from cell-free expression of DynA (left-most lane) alongside purified DynA standards at increasing concentrations (middle lanes) and molecular weight marker (right-most lane). The gel was post-stained with Coomassie stain. b) The integrated area under each band intensity peak, measured across the full width of the peak from end to end, was plotted against the corresponding amounts of purified DynA to generate a standard calibration curve. Linear regression of the calibration data was used to determine the amount of DynA produced in the PURE reaction, yielding 2.95 µmol total DynA, corresponding to a final concentration of 1.1 µM in the PURE mixture. 10 ng/µL of DynA template was used, temperature was 37 °C.

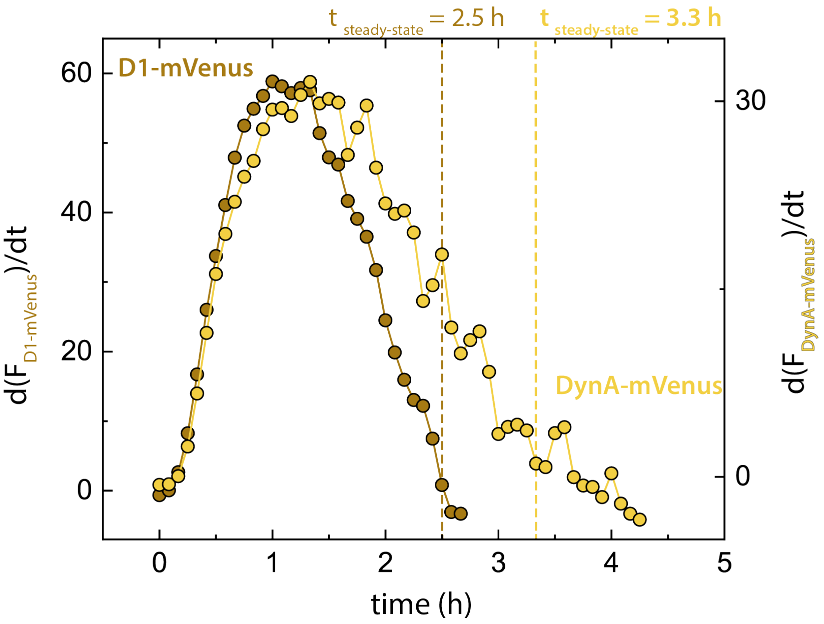


**Fig. S3.** Fluorescence traces from Figure 1d (main text) and Figure 5c (main text) were differentiated with respect to time (dF/dt), revealing that steady-states for DynA-mVenus and D1-mVenus expression are achieved at 3.3 h and 2.5 h respectively.

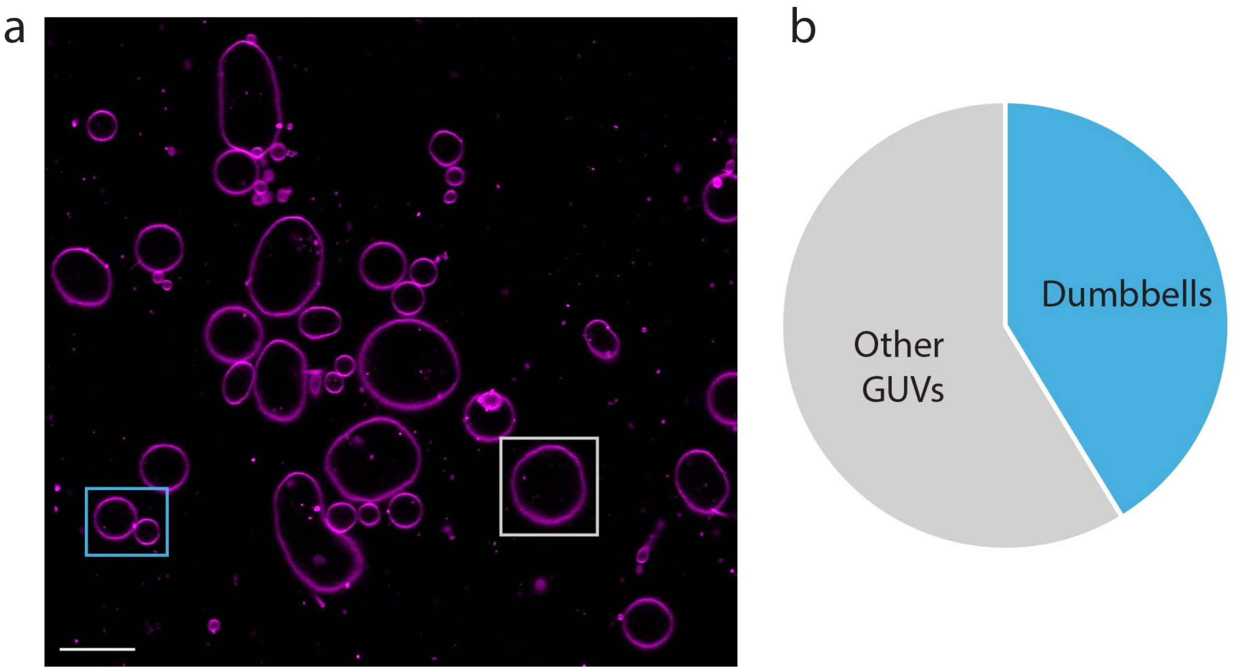


**Fig. S4.** GUVs form dumbbells when different sugars are used in the inner and outer medium while preparing the GUVs. a) Representative confocal image of GUVs prepared with 8% DOPG lipids, using 100 mM sucrose in the inner solution (IAS) and 100 mM glucose in the outer solution (OAS). A pre-deformed dumbbell-shaped GUV is highlighted in the blue box, and other GUVs are highlighted in the grey box. The osmolality ratio between outer and inner aqueous solutions were maintained to be 0.9. Scale bar: 20 µm. b) Pie chart showing the proportion of dumbbell-shaped and other GUVs observed after 1 hour of preparation. A total of 514 GUVs were analyzed.

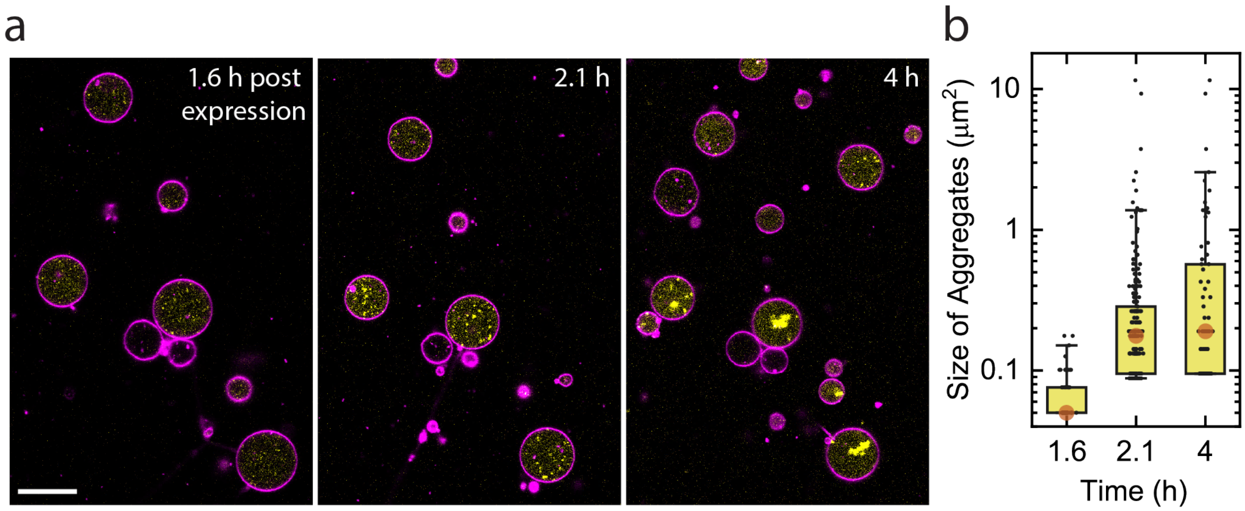


**Fig S5.** DynA-mVenus expressed within GUVs prepared with 8% DOPG forms aggregates that progressively increase in size over time. a) Time-series showing cell-free expression of DynA-mVenus over a period of 4 hours. GUVs were prepared with 8% DOPG using 100 mM sucrose in the IAS and 100 mM glucose in the OAS, with 20 ng/µL DynA-mVenus template and incubated at 37 °C. Scale bar: 20 µm. b) Quantification of DynA-mVenus aggregate size at each time point, where aggregate area was used as a proxy for aggregate size (n = 54, 194, and 70, respectively). Each data point represents an individual aggregate. In the box plots, boxes indicate 25–75% of data, whiskers represent 5–95% of data, and the solid orange circle denotes the median aggregate size.

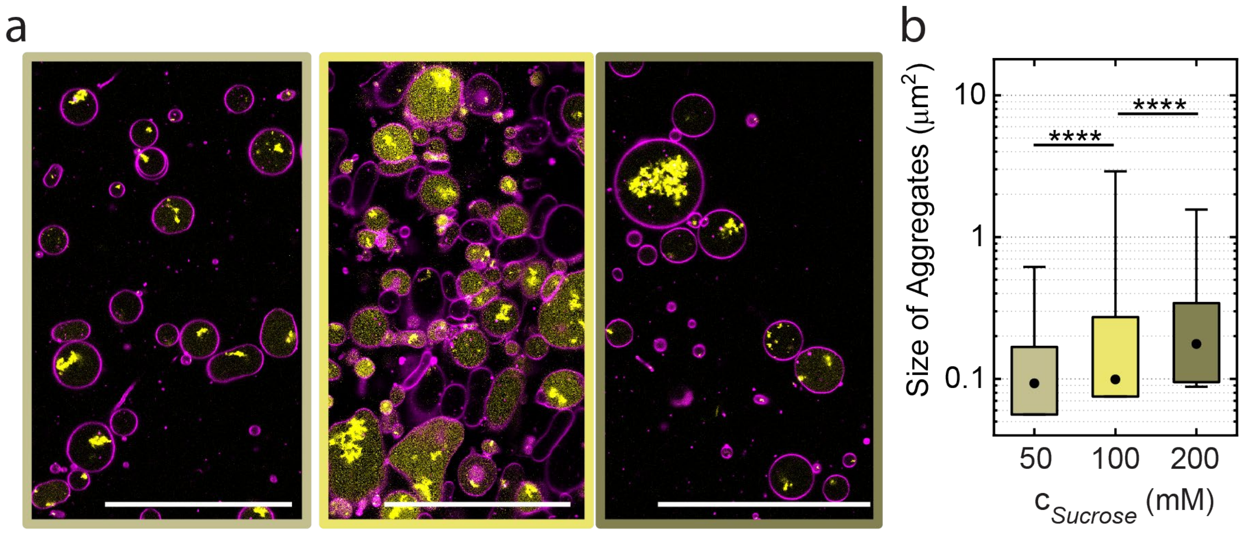


**Fig S6.** Effect of sucrose concentration on the aggregation of cell-free expressed DynA-mVenus within GUVs prepared using 8% DOPG. a) Representative images of DynA-mVenus expressed within GUVs after 4 hours of cell-free expression under increasing sucrose concentrations in the IAS from left to right: 50 mM, 100 mM, and 200 mM. Corresponding glucose concentrations were used in the OAS. GUVs were prepared using 8% DOPG with 20 ng/µL DynA-mVenus template and incubated at 37 °C. Scale bar: 100 µm. b) Quantification of DynA-mVenus aggregate size under increasing sucrose concentrations, where aggregate area was used as a proxy for aggregate size (n = 3204, 6884, and 264, respectively). In the box plots, boxes indicate 25–75% of data, whiskers represent 5–95% of data, and the solid black circle denotes the median aggregate size. Statistical significance was tested using a non-parametric Mann-Whitney test (**** $p<0.0001$)

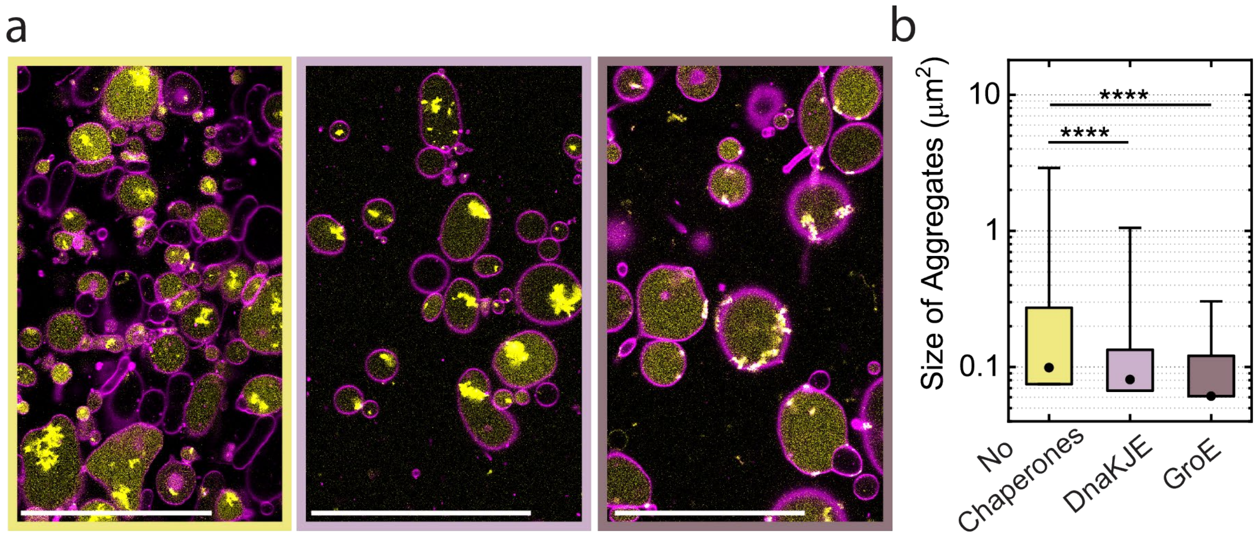


**Fig. S7**. Effect of supplementing chaperones on the aggregation of cell-free expressed DynA-mVenus within GUVs prepared using 8% DOPG. a) Representative images of DynA-mVenus expressed within GUVs after 4 hours of cell-free expression performed either in the absence or presence of different chaperone systems. From left to right: standard conditions without chaperones (20 ng/µL plasmid DNA; yellow), supplementing with DnaKJE mix (1 µM; light purple), or GroE mix (0.5 µM GroEL and 1 µM GroES; dark purple). GUVs were prepared with 8% DOPG using 100 mM sucrose in the IAS and 100 mM glucose in the OAS and incubated at 37 °C. Scale bar: 100 µm. b) Quantification of DynA-mVenus aggregate size under the three corresponding chaperone conditions, where aggregate area was used as a proxy for aggregate size (n = 6884, 2226, and 6087, respectively). In the box plots, boxes indicate 25–75% of data, whiskers represent 5–95% of data, and the solid black circle denotes the median aggregate size. Statistical significance was tested using a non-parametric Mann-Whitney test (**** $p<0.0001$).

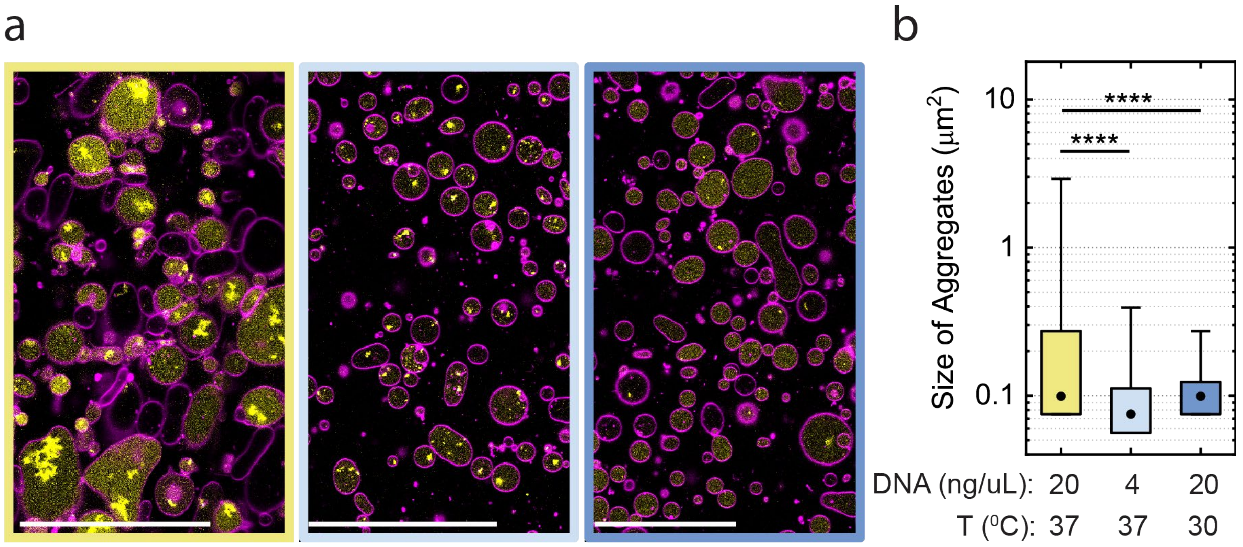


**Fig. S8.** Effect of the environmental conditions on the aggregation of cell-free expressed DynA-mVenus within GUVs prepared using 8% DOPG. a) Representative images of DynA-mVenus expressed within GUVs after 4 hours of cell-free expression under three environmental conditions. From left to right: standard conditions (20 ng/μL plasmid DNA, 37 °C; yellow), reduced plasmid concentration (4 ng/μL plasmid DNA; light blue), and reduced incubation temperature (30 °C with 20 ng/μL plasmid DNA; dark blue). GUVs were prepared with 8% DOPG using 100 mM sucrose in the IAS and 100 mM glucose in the OAS. Scale bar: 100 μm. b) Quantification of DynA-mVenus aggregate size under the three corresponding environmental conditions, where aggregate area was used as a proxy for aggregate size (n = 6884, 71761, and 60876, respectively). In the box plots, boxes indicate 25–75% of data, whiskers represent 5–95% of data, and the solid black circle denotes the median aggregate size. Statistical significance was tested using a non-parametric Mann-Whitney test (**** $p<0.0001$).

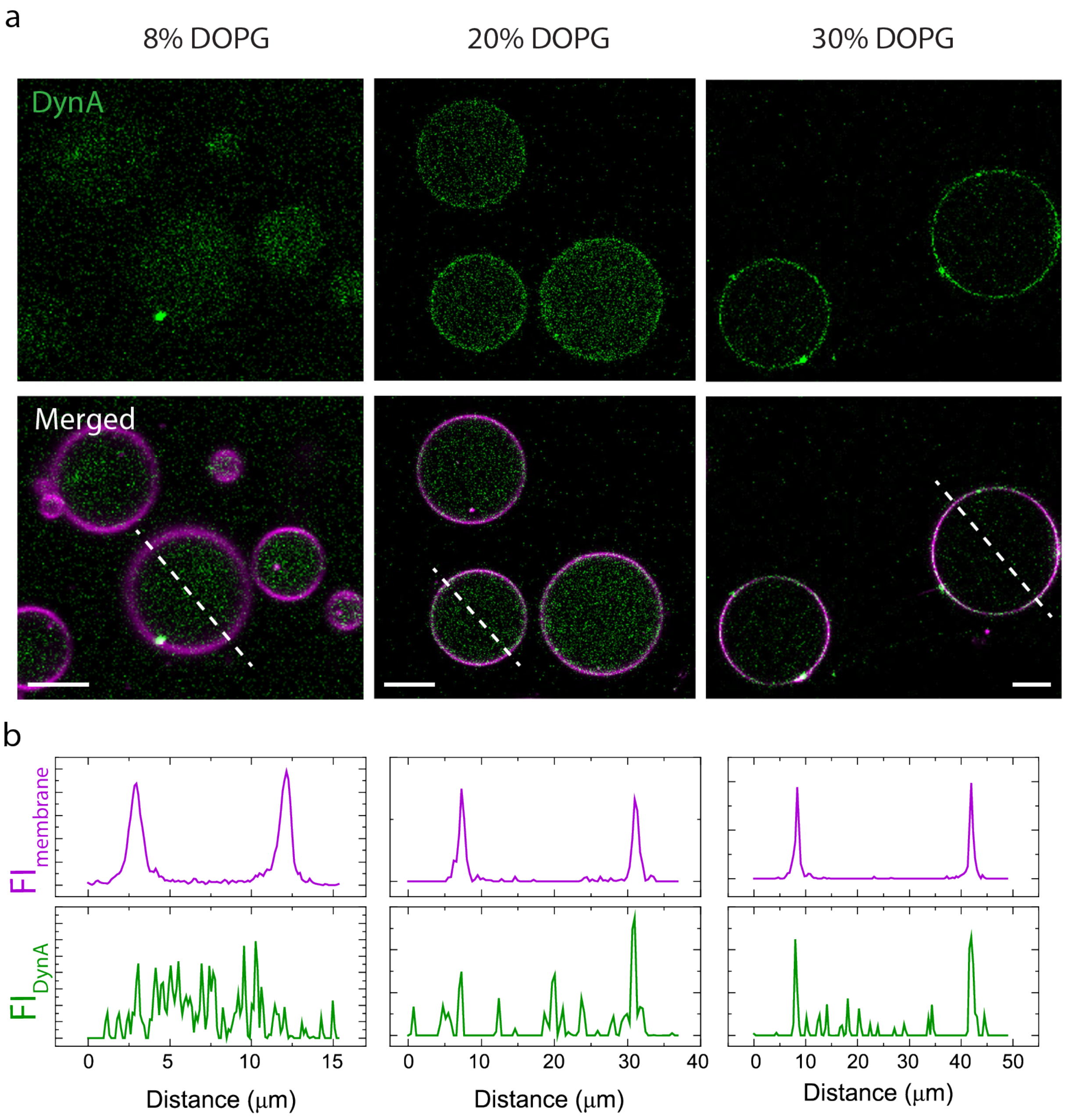


**Fig. S9**. DynA preferentially binds to the membranes of GUVs prepared using higher DOPG concentrations. a) Representative 2D confocal images of the GUVs prepared using increasing DOPG concentrations (8%, 20%, and 30%). GUVs were prepared using IAS composed of 37% Optiprep in 50 mM Tris-HCl buffer supplemented with 5 mM $MgCl_2$ and an OAS consisting of 50 mM Tris-HCl buffer with glucose added to maintain an osmolality ratio of 0.85 relative to IAS. 300 nM purified DynA was used in all conditions. Scale bars: 5 µm (left panel), 10 µm (middle panel), and 10 µm (right panel). b) Line-segment analysis shows that DynA resides within the GUV lumen for lower DOPG concentration whereas enriches at the membrane for DOPG concentrations ≥ 20%.

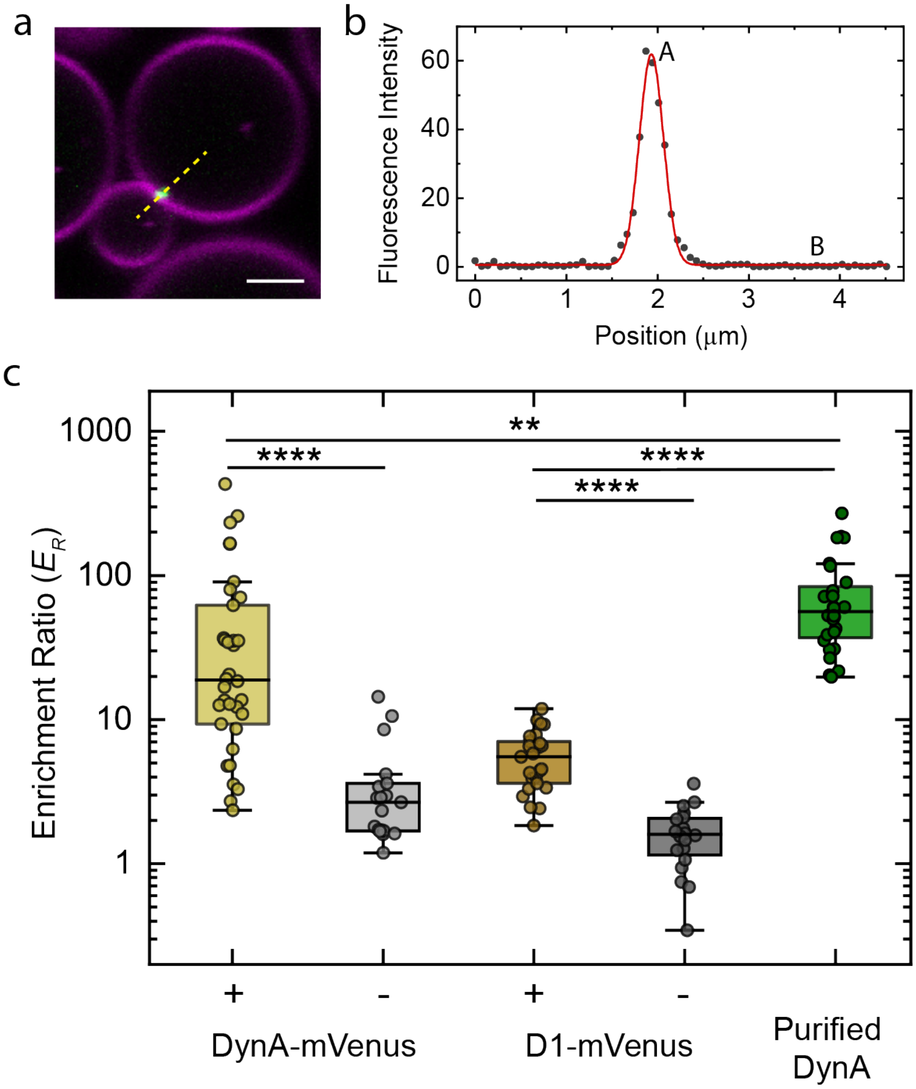


**Fig. S10.** Estimating the enrichment ratio of cell-free expressed DynA-mVenus and D1-mVenus at the neck of dumbbells. a) Fluorescence intensity line profiles were extracted across the dumbbell neck. b) The line profile is fitted with a Gaussian Amplitude function to extract peak fluorescence intensity, A and the luminal fluorescence intensity, B. The ratio between A and B is called enrichment ratio ($E_R$). c) Comparison between the enrichment ratio of cell-free expressed DynA-mVenus and D1-mVenus with purified DynA at the necks of dumbbell GUVs. For cell-free expressed proteins, the dumbbells with visually prevalent protein enrichment are compared with dumbbells where no enrichment can be observed. For each condition, n = 34, 19, 25, 20, and 28 dumbbells (N =2) were analyzed. In the box plots, boxes indicate 25–75% of data, whiskers represent 5–95% of data, and the solid black circle denotes the median aggregate size. Statistical significance was tested using a non-parametric Mann-Whitney test.

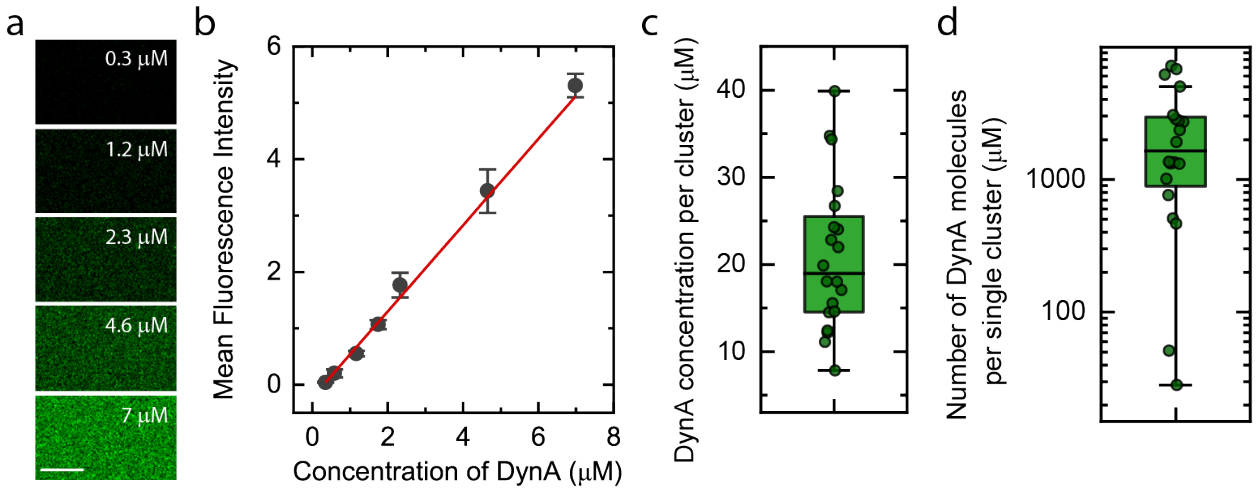


**Fig. S11.** Estimating DynA concentration within the clusters formed at the neck of the dumbbells. a) For quantitative imaging, we first performed an xz-scan to locate the precise z-position of the coverslip. We then acquired xy-confocal images of DynA solutions at varying concentrations in photon counting mode while maintaining the z-position between 12-15 μm. Scale bar: 2.5 μm. b) To determine the unknown DynA concentrations within the DynA clusters at the neck of dumbbells, we constructed a concentration-intensity calibration curve. The mean pixel value over the entire 512 × 512-pixel image was calculated for five different images per condition and used as the mean fluorescence intensity for that DynA concentration, represented by the solid grey data points. The data points were fitted with a linear model, represented by red line, yielding a slope and intercept of 0.765 and -0.228, respectively. The concentration (c) and number (d) of DynA molecules present within a single DynA cluster was then estimated by converting the measured mean fluorescence intensity of each cluster into concentration using the calibration curve constructed in (b).

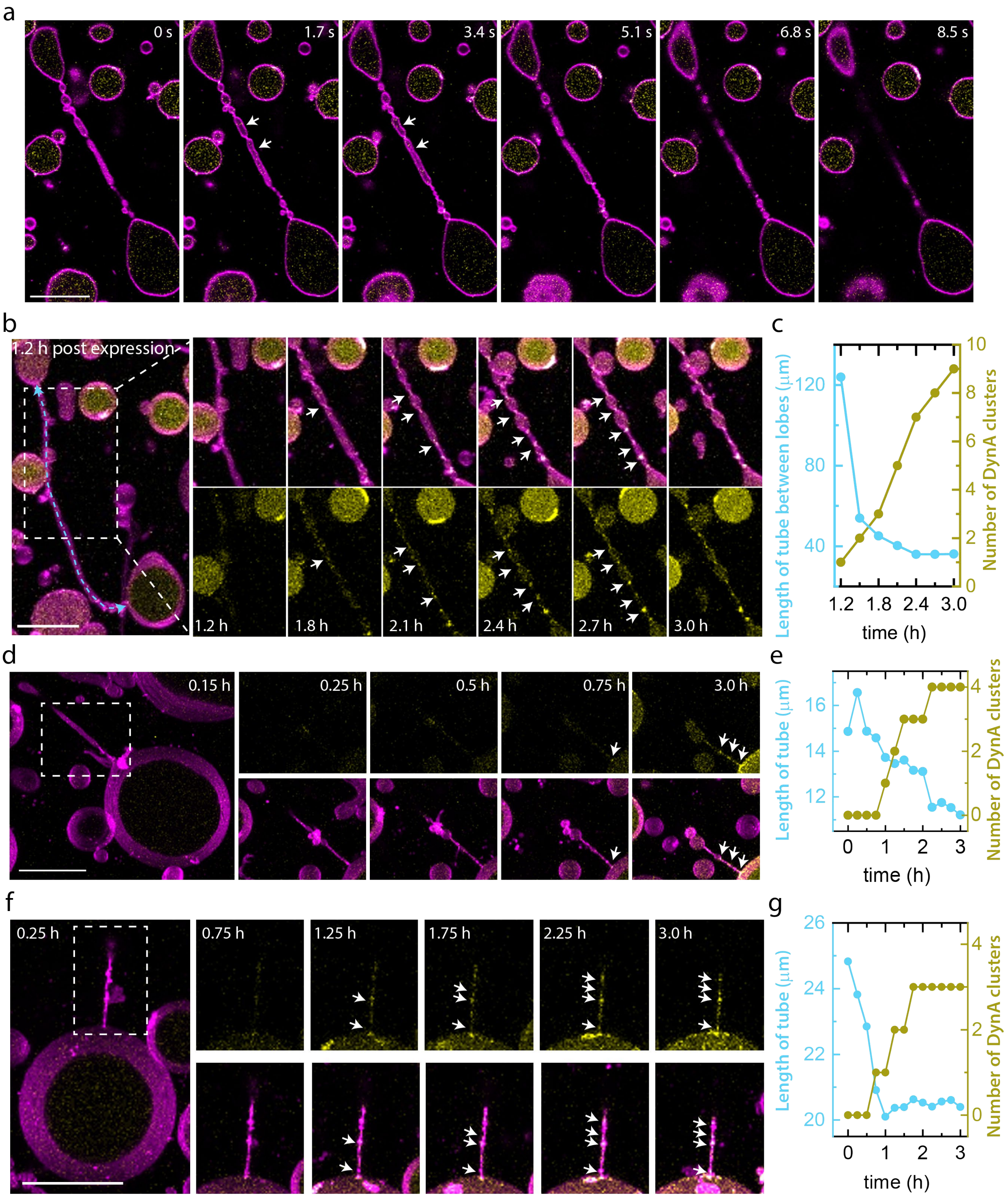
a
0 s
1.7 s
3.4 s
5.1 s
6.8 s
8.5 s
b
1.2 h post expression
1.2 h
1.8 h
2.1 h
2.4 h
2.7 h
3.0 h
c
Length of tube between lobes (μm)
Number of DynA clusters
time (h)
d
0.15 h
0.25 h
0.5 h
0.75 h
3.0 h
e
Length of tube (μm)
Number of DynA clusters
time (h)
f
0.25 h
0.75 h
1.25 h
1.75 h
2.25 h
3.0 h
g
Length of tube (μm)
Number of DynA clusters
time (h)

**Fig. S12**. DynA stabilizes regions of high curvature within a thermally fluctuating cylindrical tube, transforming it into a chain of dumbbells. a) xy scans of a wide cylindrical tube at different z positions show the tube undergoing thermal fluctuations, creating transient negatively curved sites marked by white arrows. b, d, f) 3D confocal z-stack projections of three distinct pre-deformed GUVs show a cylindrical tube transforming into a chain of pearls with multiple lobes, where each emerging neck becomes enriched with DynA-mVenus as expression progresses over 3 hours. c, e, g) The respective longitudinal contraction of the tube coincides with the time-dependent appearance of DynA-mVenus-localized necks, indicating DynA driven stabilization of highly curved necks, resulting in a constricted neck topology. Scale bars: 20 µm.

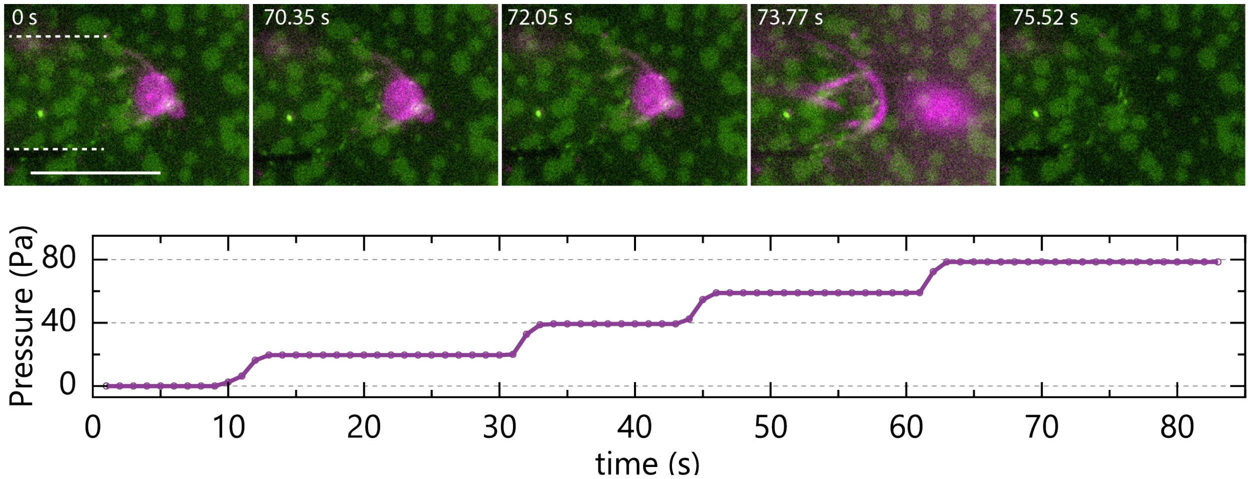


**Fig. S13**. Representative trace of pressure versus time for a low-pressure regime micropipette aspiration set-up with critical aspiration pressure, $P_C$ of 78 Pa. The pressure trace is supplemented with representative fluorescence microscopic images over time. Scale bar: 10 µm. Membrane is shown in magenta and DynA in green. The out of focus micropipette is marked in white dashed line.

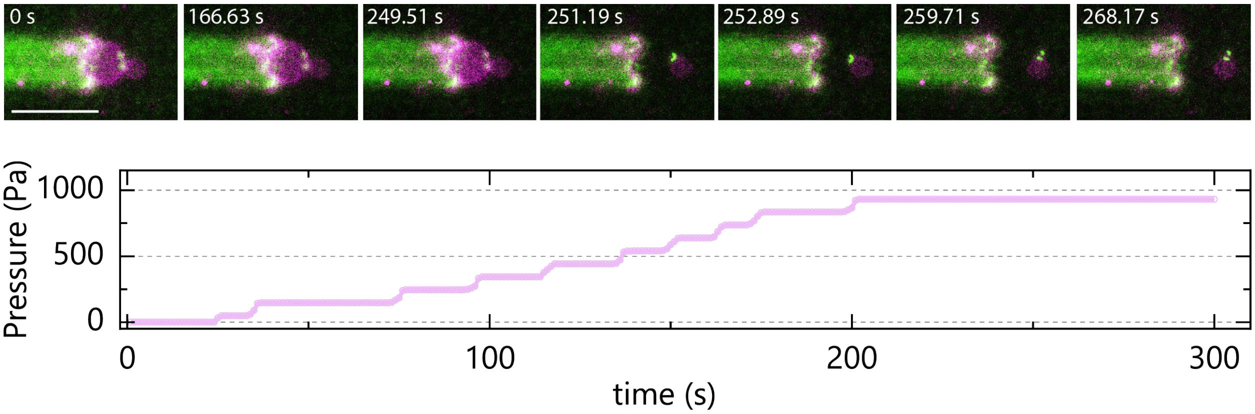


**Fig. S14**. Representative trace of pressure versus time for a high-pressure regime micropipette aspiration set-up with $P_C$ of 932 Pa. The pressure trace is supplemented with representative fluorescence microscopic images over time. Scale bar: 10 µm. Membrane is shown in magenta and DynA in green. Due to autofluorescence from the BSA coating, the micropipette is also visible in the green channel.

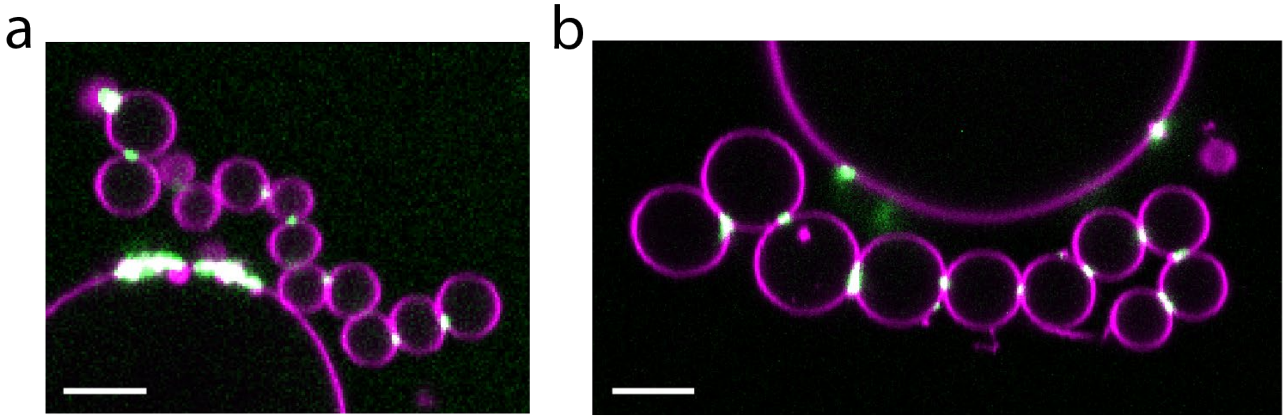


**Fig. S15**. GUVs remained adhered after $Mg^{2+}$ removal from the OAS. a,b) Representative images of pre-deformed GUVs with DynA-enriched necks in the presence (a) and absence (b) of 5 mM $Mg^{2+}$ in the OAS. GUVs were prepared at an osmolality ratio of 0.92 between OAS and IAS. Purified DynA was used at 300 nM in all conditions. Scale bars: 5 µm. The membrane is shown in magenta and DynA in green.

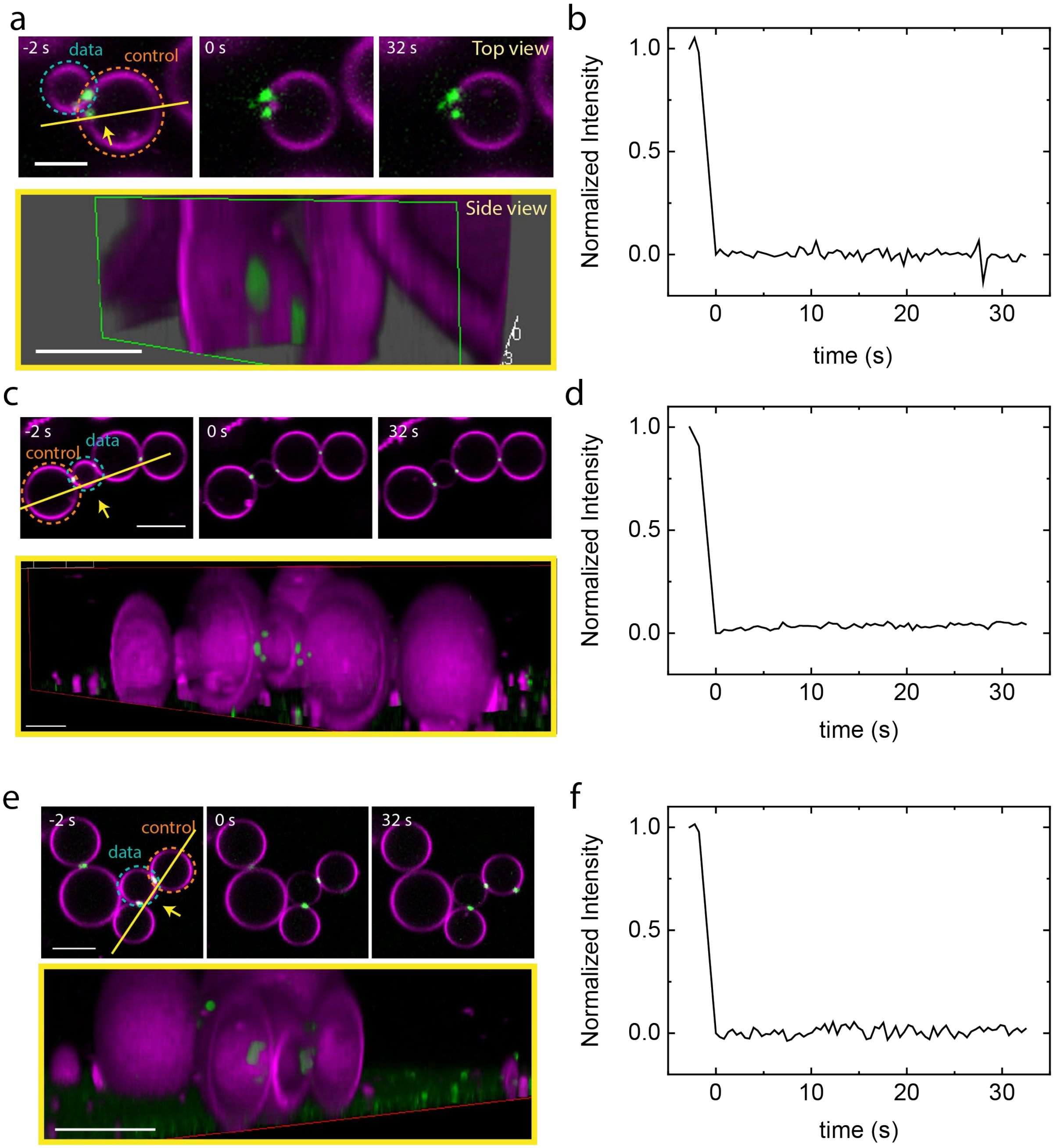


**Fig. S16**. DynA clusters at the necks of dumbbells which have undergone full scission have elongated, disc-like architectures. a), c), and e) 3D reconstructions of confocal images reveal an elongated disc-like architecture of DynA (green) enriched at the membrane neck (magenta) of 3 different dumbbells. The top panel shows the top view (xy scan) of the dumbbell, and the bottom panel shows an orthogonal optical section (side view) reconstructed from the same dataset along the yellow line, with the viewing direction indicated by the arrow. Scale bars: a) 2.5 µm (top), 2 µm (bottom); b) 2.5 µm (top), 2 µm (bottom); c) 5 µm (top and bottom). b), d), f) FRAP analysis of the membrane connectivity at the necks of dumbbells, confirm full scission.

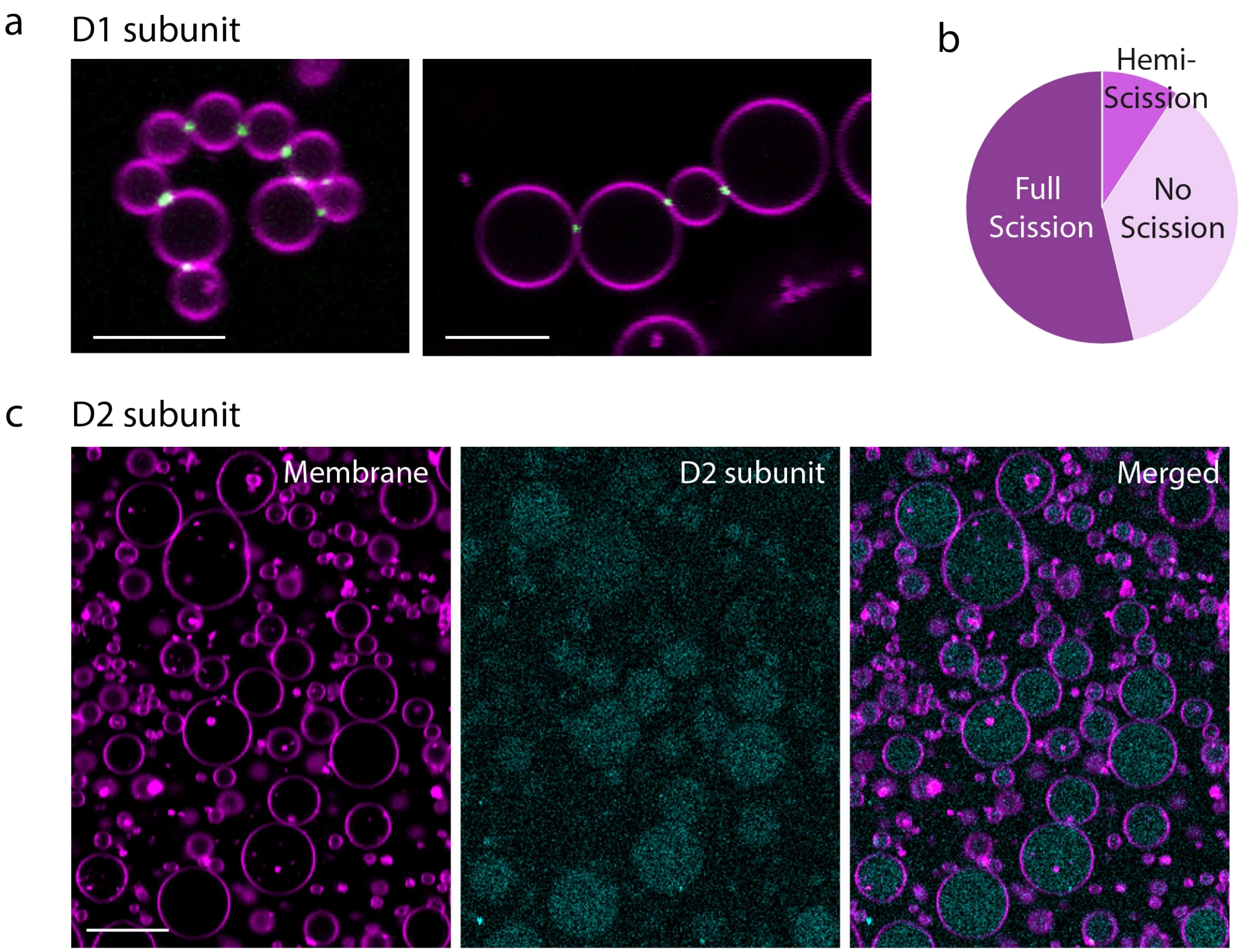


**Fig. S17**. The D1 subunit of DynA is sufficient for division of synthetic cells. a) Representative images of pre-deformed GUVs (magenta) enriched with D1 (green) at the necks. Scale bars: 5 µm. b) Pie chart indicating the fraction of full scission, hemi-scission and no scission events at the necks of dumbbells enriched with D1 as determined by FRAP assays of lipid diffusion across necks (n = 54 from 2 independent preparations). c) D2 subunit resides within the lumen and does not enrich at the necks of dumbbells. Scale bars: 10 µm. GUVs were prepared using IAS composed of 37% Optiprep in 50 mM Tris-HCl buffer supplemented with 5 mM $MgCl_2$ and an OAS consisting of 50 mM Tris-HCl buffer with glucose added to maintain an osmolality ratio of 0.92 relative to IAS. Purified D1 and D2 were used at 300 nM in all conditions.

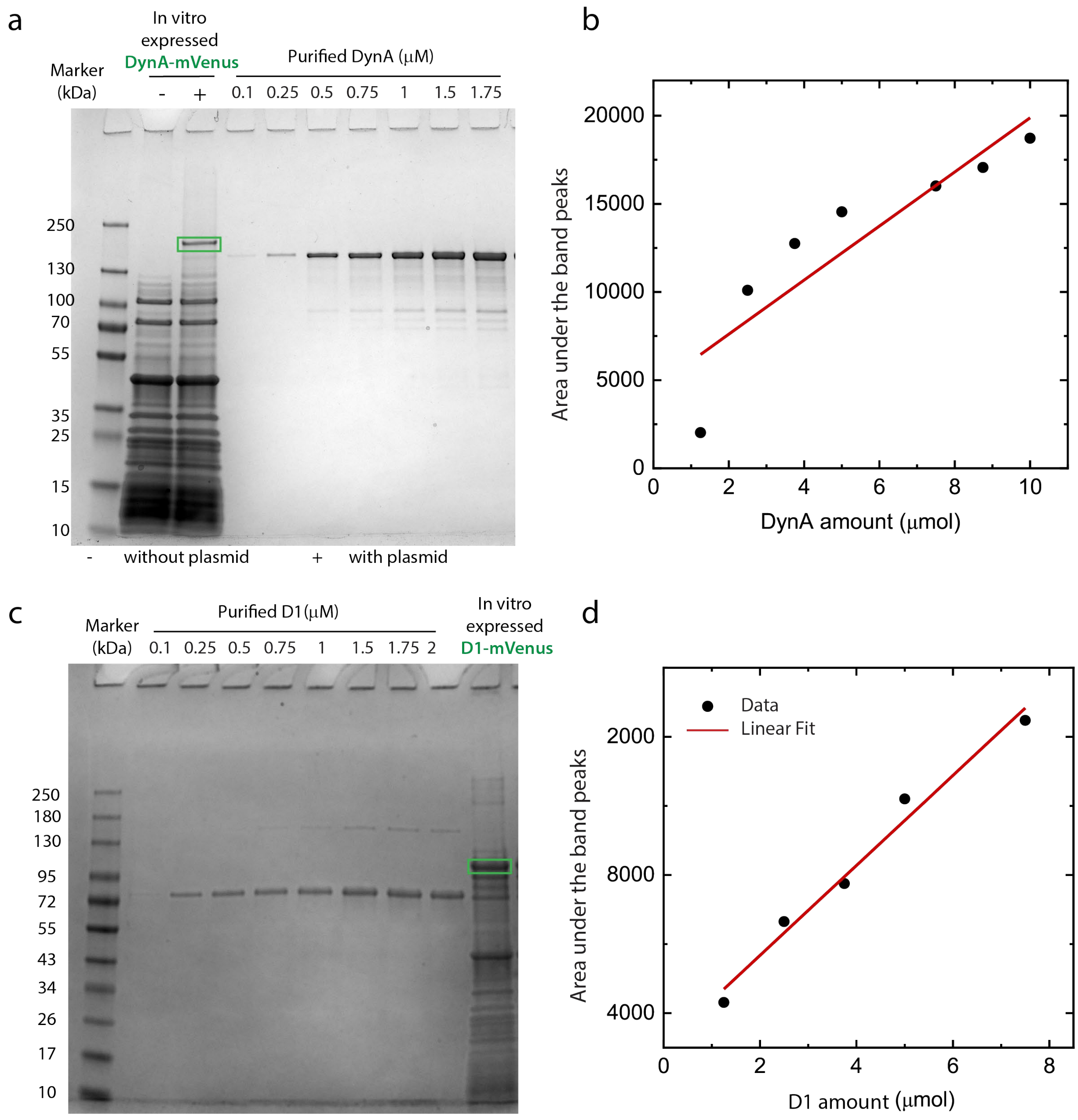


**Fig. S18**. Measurement of cell-free expressed DynA-mVenus and D1-mVenus concentrations using the PURE system. a and c) SDS-PAGE analysis of translated products obtained from cell-free expression of DynA-mVenus (a) and D1-mVenus (c) (bands indicated in green boxes) alongside increasing concentrations of purified DynA and D1 standards, respectively. Gels were post-stained with Coomassie stain. The gel also shows a control measurement of the PURE solution without added DNA template, showing the DynA-mVenus band indeed results from expression. b and d) The integrated areas under each band intensity peak, measured across the full width of the peak from end to end, were plotted against the corresponding amounts of purified DynA (b) and D1 (d) to generate protein-specific calibration curves. Linear regression of these standards was used to estimate the amounts of fusion proteins produced in the PURE reactions. This analysis yielded an apparent production of 2.1 µmol DynA–mVenus (0.43 µM) and 6.35 µmol D1–mVenus (2.7 µM) from 10 uL volumes at 37 °C using DNA templates at 10 ng/µL for both constructs. The reported values represent uncorrected densitometric estimates based on calibration with non-fused purified proteins and were not corrected for differences in molecular weight or Coomassie dye-binding properties introduced by the mVenus fusion tag. Overall, D1–mVenus was expressed at approximately six-fold higher yield than DynA–mVenus under the conditions tested.

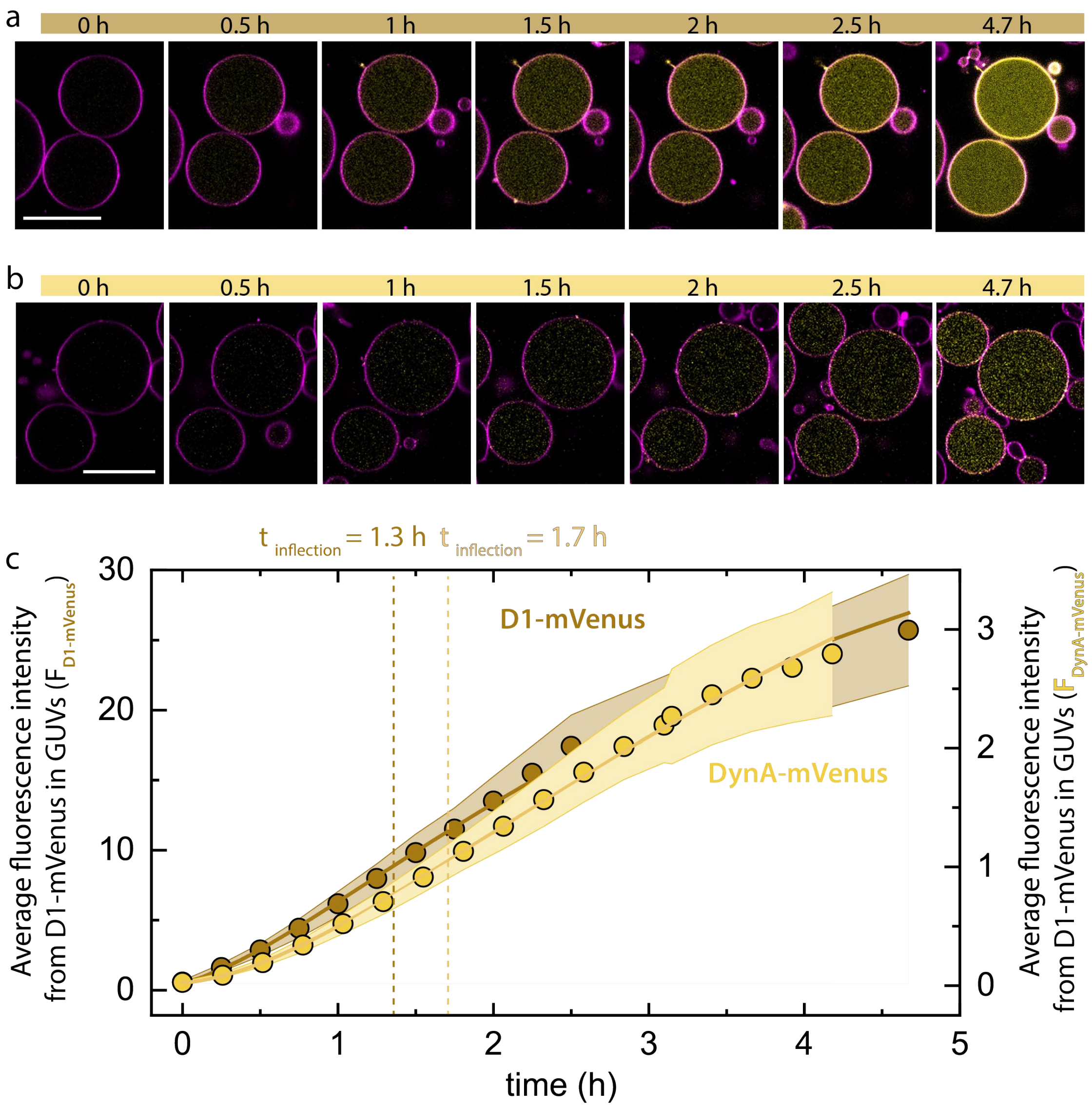


**Fig. S19**. Time-dependent fluorescence intensity of D1-mVenus and DynA-mVenus during cell-free expression in GUVs. a, b) Representative time-lapse fluorescence images showing the cell-free expression of D1-mVenus (a) and DynA-mVenus (b) within GUVs. Images were taken at the equatorial plane of the GUVs. c) Average of mean fluorescence intensity from D1-mVenus and DynA-mVenus within GUVs is plotted over time. Data points represent the mean fluorescence intensity averaged across 24 individual GUVs for each construct, while the shaded regions indicate the standard deviation. To compare the kinetics of protein accumulation, the data is fitted with standard Gompertz growth model (solid lines) to determine the inflection points for DynA-mVenus and D1-mVenus. D1–mVenus reached its maximum accumulation rate at approximately 1.3 h (inflection point), after which the rate declined, indicating the onset of saturation. In contrast, DynA–mVenus reached its maximum accumulation rate at approximately 1.7 h. Collectively, these results indicate that D1–mVenus exhibits both faster expression kinetics and an earlier approach to steady-state accumulation than DynA–mVenus.

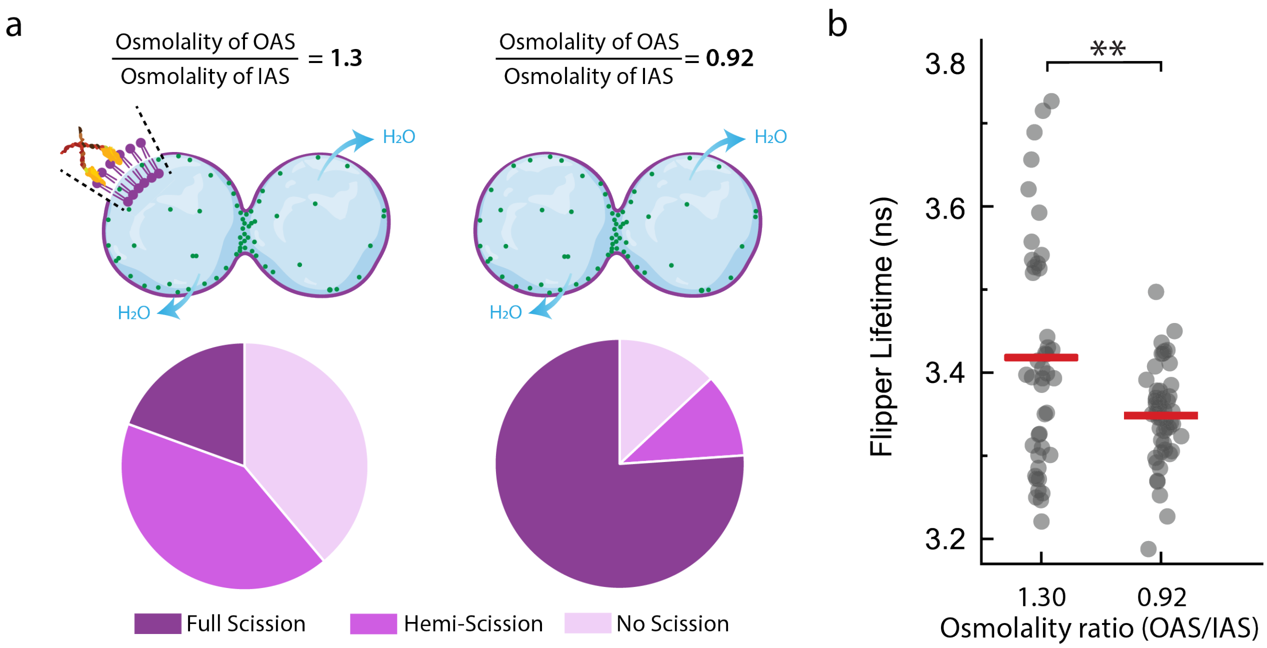


**Fig. S20**. Comparison of membrane-remodeling events induced by DynA in membrane dumbbells prepared under different osmotic conditions. a) Pie charts indicating the percentage of full scission, hemi-scission and no scission events at the necks of dumbbells enriched with purified DynA, compared under two preparation conditions. The left panel shows the condition reported previously (data adapted from de Franceschi et al., 2023), in which DNA nanostars were added to OAS, maintaining an osmolality ratio of 1.3 between OAS and IAS. The right panel shows the optimized condition in our study, in which an osmolality ratio of 0.92 was used (n = 46 from 2 independent preparations). Dumbbells were prepared as specified in Table 3, except that 250 nM of DNA nanostars and 5 mM $MgCl_2$ were included in the OAS for condition shown in the left panel. Purified DynA was used at 300 nM in both conditions. b) Mean intensity weighted fluorescence lifetimes of FliptR measured within dumbbells prepared under the two conditions specified in (a). Each data point represents the average lifetime value of one individual GUV (n = 43 and 54 GUVs for left and right condition respectively). Red bars indicate the mean lifetime for each dataset. Statistical significance was tested using two sample t-test after Welch correction (**$p < 0.01$).

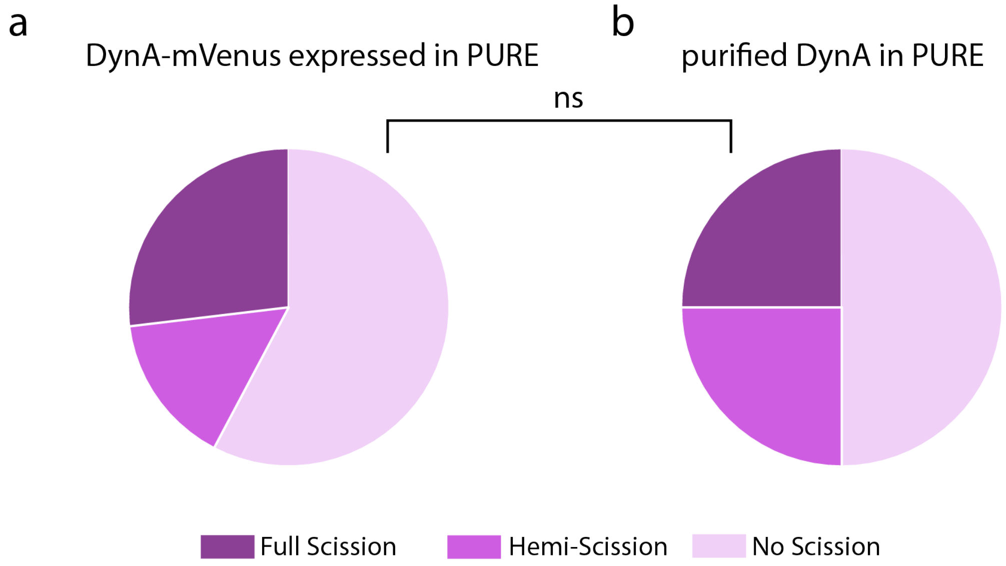


**Fig. S21.** Comparison of membrane-remodeling events induced by DynA-mVenus produced by cell-free expression and purified DynA. a) Membrane remodeling events as quantified by FRAP assays induced by DynA-mVenus synthesized in situ using the PURE cell-free expression system. b) Membrane remodeling events induced by purified DynA encapsulated together with all the components of the cell-free expression system used in (a), except for the DNA template encoding DynA-mVenus. Data obtained from n = 26, across 3 and 2 independent preparations for each condition respectively. The two conditions were statistically non-significant from each other when tested using two-sample t-test after Welch correction ($p>0.05$).

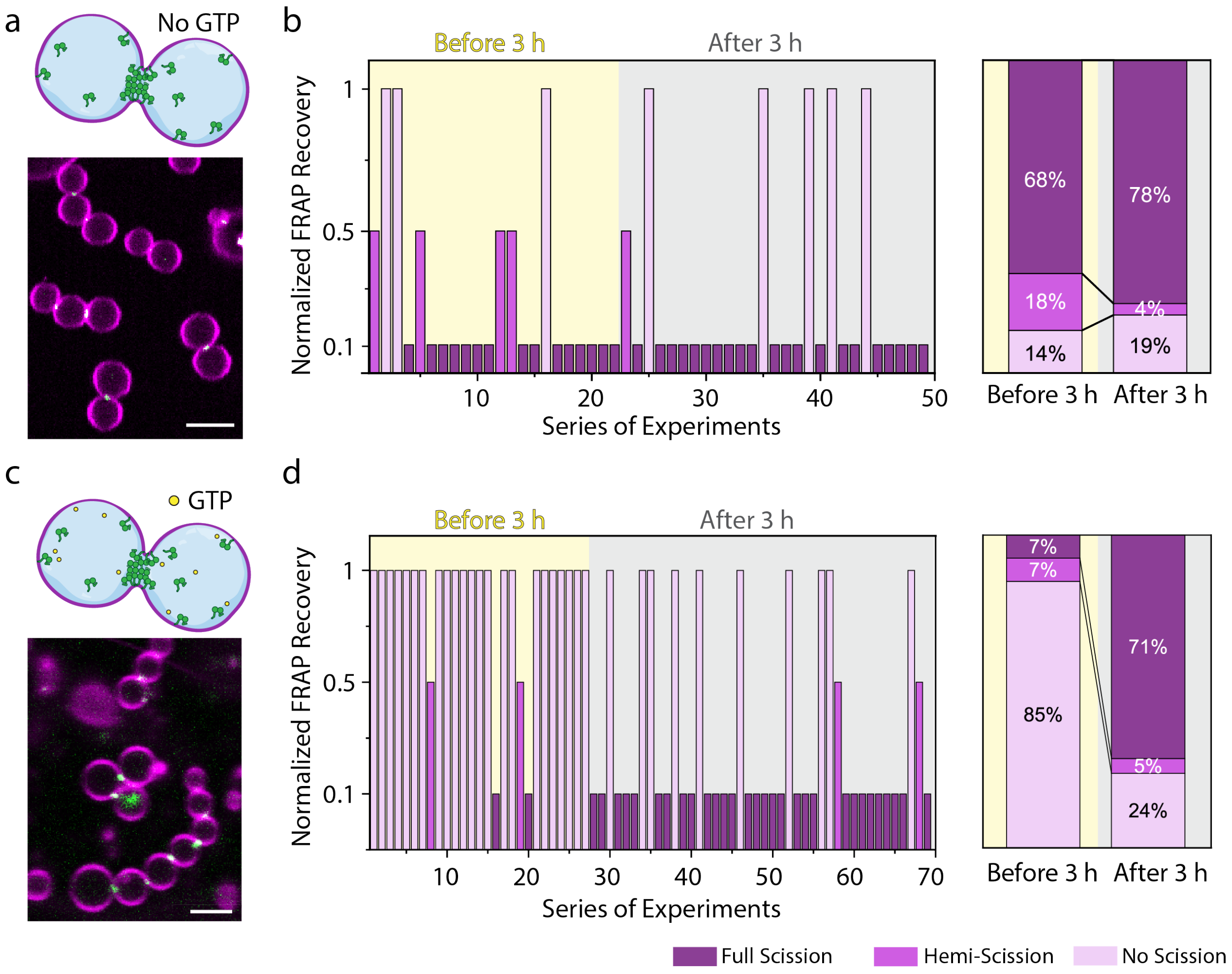


**Fig. S22.** DynA loses its scission activity in the presence of GTP which is recovered after 3 h, likely reflecting the GTP hydrolysis cycle of DynA. a and c) Representative images of pre-deformed GUVs with DynA-enriched necks in the absence (a) and presence (c) of 2 mM GTP in the IAS. b and d) Left panels show the normalized FRAP recovery for each experiment in the order they were performed. A normalized recovery value of 1 indicates an open neck, 0.5 indicates hemi-scission, and 0.1 indicates complete scission (a value of 0.1 was assigned instead of 0 to facilitate data visualization). Right panels show the normalized FRAP results grouped according to the membrane remodeling events observed before and after 3 h. GUVs were prepared with an OAS/IAS osmolality ratio of 0.92. Purified DynA was used at a concentration of 300 nM under both conditions. Scale bars: 2.5 µm.

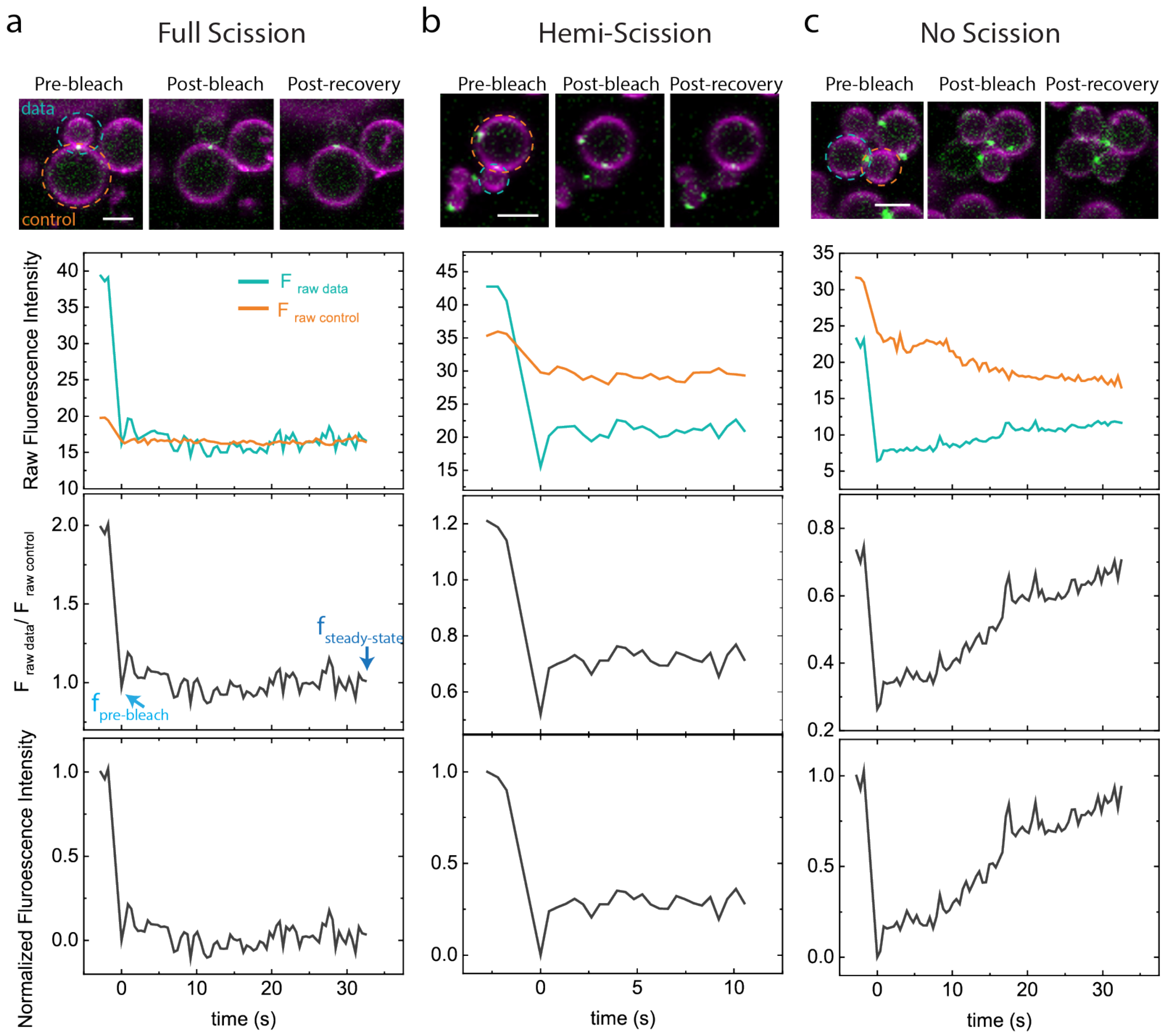


**Fig. S23**. Quantification of fluorescence recovery in FRAP experiments for the classification of full scission, hemi-scission, and no-scission events. a, b, and c) Representative examples of full-scission, hemi-scission, and no-scission events are shown. Top panel: Representative confocal images acquired during the pre-bleach, post-bleach, and post-recovery stages. The bleached lobe of the dumbbell is outlined in blue and designated as the data lobe, while the connected, unbleached lobe is outlined in orange and serves as the control lobe. Middle panel: Raw fluorescence intensity traces for the bleached (data) and control lobes. The fluorescence intensity of the bleached lobe is plotted relative to that of the control lobe to correct for imaging-related intensity fluctuations. From these traces, the fluorescence intensity before bleaching ($f_{pre\text{-}bleach}$) and the steady-state fluorescence intensity after recovery ($f_{steady\text{-}state}$) are determined. These parameters are subsequently used to normalize the fluorescence recovery of the bleached lobe relative to the control lobe.
